# A missense mutation dissociates triglyceride and phospholipid transfer activities in zebrafish and human microsomal triglyceride transfer protein

**DOI:** 10.1101/701813

**Authors:** Meredith H. Wilson, Sujith Rajan, Aidan Danoff, Richard J. White, Monica R. Hensley, Vanessa H. Quinlivan, James H. Thierer, Elisabeth M. Busch-Nentwich, M. Mahmood Hussain, Steven A. Farber

## Abstract

Microsomal triglyceride transfer protein (MTP) transfers triglycerides and phospholipids and is essential for the assembly of Apolipoprotein B (ApoB)-containing lipoproteins in the endoplasmic reticulum. We have discovered a zebrafish mutant (*mttp^c655^*) expressing a C-terminal missense mutation (G863V) in Mttp, one of the two subunits of MTP, that is defective at transferring triglycerides, but retains phospholipid transfer activity. Mutagenesis of the conserved glycine in the human MTTP protein (G865V) also eliminates triglyceride but not phospholipid transfer activity. The G863V mutation reduces the production and size of ApoB-containing lipoproteins in zebrafish embryos and results in the accumulation of cytoplasmic lipid droplets in the yolk syncytial layer. However, *mttp^c655^* mutants exhibit only mild intestinal lipid malabsorption and normal growth as adults. In contrast, zebrafish mutants bearing the previously identified *mttp^stl^* mutation (L475P) are deficient in transferring both triglycerides and phospholipids and exhibit gross intestinal lipid accumulation and defective growth. Thus, the G863V point mutation provides the first evidence that the triglyceride and phospholipid transfer functions of a vertebrate MTP protein can be separated, arguing that selective inhibition of the triglyceride transfer activity of MTP may be a feasible therapeutic approach for dyslipidemia.

## INTRODUCTION

In vertebrates, Apolipoprotein B-containing lipoproteins (B-lps) are produced by the intestine and liver and transport lipid and fat-soluble vitamins to the peripheral tissues through the circulation. Lipoproteins are composed of a neutral core of triglyceride (TG) and cholesteryl esters surrounded by a monolayer of phospholipid, free cholesterol and sphingomyelin. B-lps contain one apolipoprotein B (ApoB) scaffold protein embedded in the phospholipid monolayer as well as other exchangeable lipoproteins (Hussain et al., 1996; Schumaker et al., 1994). B-lp assembly occurs in the endoplasmic reticulum (ER) and requires the activity of microsomal triglyceride transfer protein (MTP) (Hussain et al., 2003b; Patel and Grundy, 1996; Wetterau and Zilversmit, 1986). As ApoB is translated and translocated into the lumen of the ER, MTP physically interacts with and transfers lipids to ApoB to form primordial lipoproteins (Bradbury et al., 1999; Hussain et al., 2003b; Patel and Grundy, 1996; Wu et al., 1996). MTP may also aid in the formation of TG-rich lumenal lipid droplets that are believed to fuse to the primordial lipoproteins, thus increasing the size of nascent lipoproteins (Alexander et al., 1976; Boren et al., 1994; Hamilton et al., 1998; Kulinski et al., 2002; Raabe et al., 1999; Wang et al., 1997).

MTP is a heterodimer of the large 97-kDa M subunit (microsomal triglyceride transfer protein, MTTP) and the small 58-kDa P subunit protein disulfide isomerase (PDI)(Wetterau et al., 1990). Mutations in the *MTTP* gene that prevent lipid transfer and ApoB secretion cause the disease Abetalipoproteinemia (OMIM 200100), characterized by a virtual absence of plasma B-lps (Kane, 1995; Sharp et al., 1993; Shoulders et al., 1993; Wetterau et al., 1992). Patients exhibit fat malabsorption, low plasma triglyceride, and cholesterol levels as well as fat-soluble vitamin deficiencies (Kane, 1995; Lee and Hegele, 2014; Walsh and Hussain, 2016). Without adequate supplementation of essential fatty acids and fat-soluble vitamins, these patients can develop a variety of complications including neurological, opthalmological, and hematological disorders (Kane, 1995; Lee and Hegele, 2014).

Vertebrate MTP can transfer triacylglycerol, diacylglycerol, phospholipid, cholesteryl ester, ceramide, and sphingomyelin between vesicles *in vitro* (Athar et al., 2004; Iqbal et al., 2015; Jamil et al., 1995; Rava et al., 2005; Wetterau and Zilversmit, 1984, 1985). Kinetic studies suggest that MTP transiently interacts with membranes, acquires lipids, and then delivers these lipids to an acceptor membrane. The transfer of lipids occurs down a concentration gradient and does not require energy (Atzel and Wetterau, 1993, 1994). While vertebrate MTP predominantly transfers TG (Rava et al., 2005; Wetterau and Zilversmit, 1985), the Drosophila orthologue of MTP lacks TG transfer activity (Rava et al., 2006), has phospholipid transfer activity and supports secretion of vertebrate ApoB (Khatun et al., 2012; Rava and Hussain, 2007; Rava et al., 2006; Sellers et al., 2003). A further analysis of MTTP orthologues in divergent species, including nematodes, insects, fish, and mammals, indicates that all orthologues bind PDI, localize to the ER, and support human ApoB secretion (Rava and Hussain, 2007). However, only vertebrate MTP orthologues exhibit TG transfer activity, suggesting that phospholipid transfer activity was the original function of MTP orthologues and that neutral lipid transfer first evolved in fish (Rava and Hussain, 2007).

Here we describe a hypomorphic missense mutation in the C-terminal domain of zebrafish *mttp* (G863V) that decreases the production and size of B-lps *in vivo*, but has minimal effects on lipid malabsorption in the intestine and no effect on growth. Biochemical characterization of the G863V allele indicates that it is defective in triglyceride transfer activity, but retains phospholipid transfer activity. Further, we show that mutation of the conserved glycine at position 865 in human *MTTP* also selectively abolishes triglyceride transfer activity. Taken together, these data provide the first evidence that the lipid transfer functions of a vertebrate MTP can be biochemically dissociated and argues that phospholipid transfer activity is sufficient for absorption of dietary lipid.

## RESULTS

### The c655 allele is a missense mutation in mttp

During routine screening, we identified zebrafish embryos with opaque yolks (Figure 1A). In contrast to the translucent yolks of wild-type fish, when illuminated by transmitted light, the mutant yolks appear dark (Figure 1A) and, with incident light, appear off-white (Figure 1 – figure supplement 1A). The phenotype was found to be present in Mendelian ratios (Figure 1A), suggesting the presence of a homozygous recessive mutation. This new allele is identified as Carnegie allele *c655*. To map the location of the causative mutation, we did RNA sequencing of 23 opaque-yolk *c655* mutants and 23 translucent-yolk siblings and performed a Euclidean distance mapping analysis using the Mutation Mapping Analysis Pipeline for Pooled RNA-seq (MMAPPR)(Hill et al., 2013). The Loess fit to the mapping scores (Euclidean Distance^4^) (Figure 1B, top) indicated the c655 mutation was located on Chromosome 1 and lay in the region between 9-20 MB (Figure 1B, bottom). Single nucleotide variants (SNVs) present in this 11MB region in *c655* mutant embryos were assessed for their effect on annotated genes using the Ensembl Variant Effect Predictor (McLaren et al., 2016), including using the Sorting Intolerant from Tolerant algorithm (SIFT) (Sim et al., 2012), to predict the impact of changes on protein-coding sequence (tolerated or deleterious). We extracted variants that alter the protein-coding sequence as candidates for the causal mutation (223 variants in 64 genes, of which 42 are missense variants predicted to be deleterious; Supplementary File 1).

**Figure 1:**
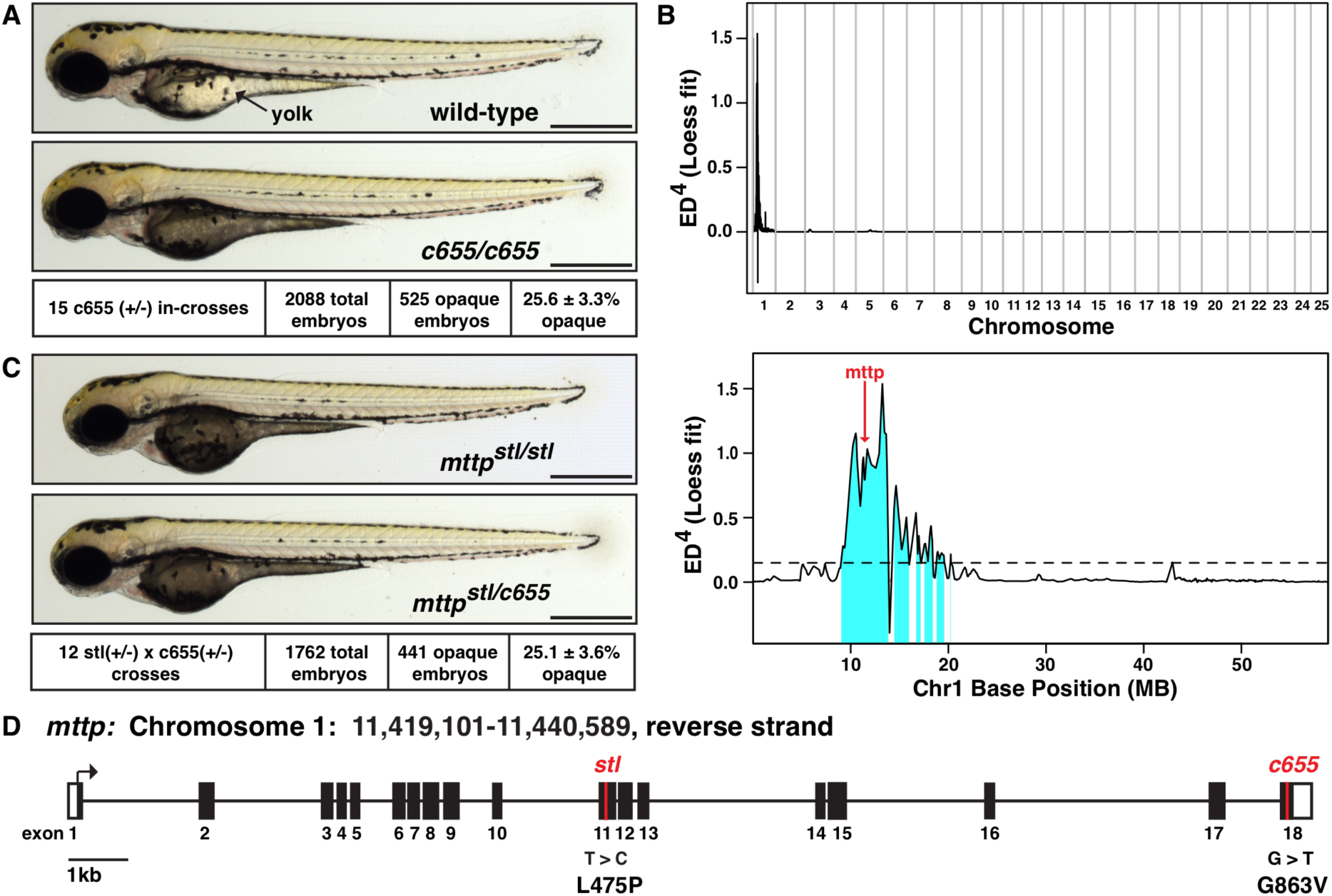
The *c655* allele is a missense mutation in the M-subunit of microsomal triglyceride transfer protein. (A) Representative images of wild-type (top) and homozygous mutant (middle) c655 zebrafish embryos at 3 days post fertilization (dpf); Scale = 500 μm. The dark/opaque yolk phenotype segregated with a Mendelian ratio consistent with a homozygous recessive mutation (bottom), mean +/-SD. (B) Euclidean distance mapping analysis plots produced by MMAPPR (Hill et al., 2013), showing the likely genomic region of the *c655* mutation: plot of the LOESS fit to the mapping scores (Euclidean Distance^4^) across all 25 chromosomes (top) and expanded view of chromosome 1(GRCz10: CM002885.1)(bottom). Red arrow shows the position of the G>T missense mutation in *mttp* (ENSDARG00000008637, position 11,421,261(GRCz10). (C) Representative images of a homozygous mutant zebrafish embryo carrying the previously described *stalactite* (*stl*) missense mutation in *mttp* (Avraham-Davidi et al., 2012)(top) and a trans-heterozygous *mttp^stl/c655^* embryo (middle); 3 dpf, scale = 500 μm. The dark/opaque yolk phenotype is present at expected ratios (bottom) and genotyping confirms only embryos with opaque yolks are trans-heterozygous for the *mttp* alleles. (D) Depiction of the *mttp* gene structure highlighting the locations of the *stl* (L475P) (position 11,431,645, GRCz10, transcript mtp-204 (ENSDART00000165753.2) nucleotide 2588) and *c655* (G863V) (nucleotide 1424) missense alleles in exon 11 and 18, respectively.

**Figure 1 – figure supplement 1:**
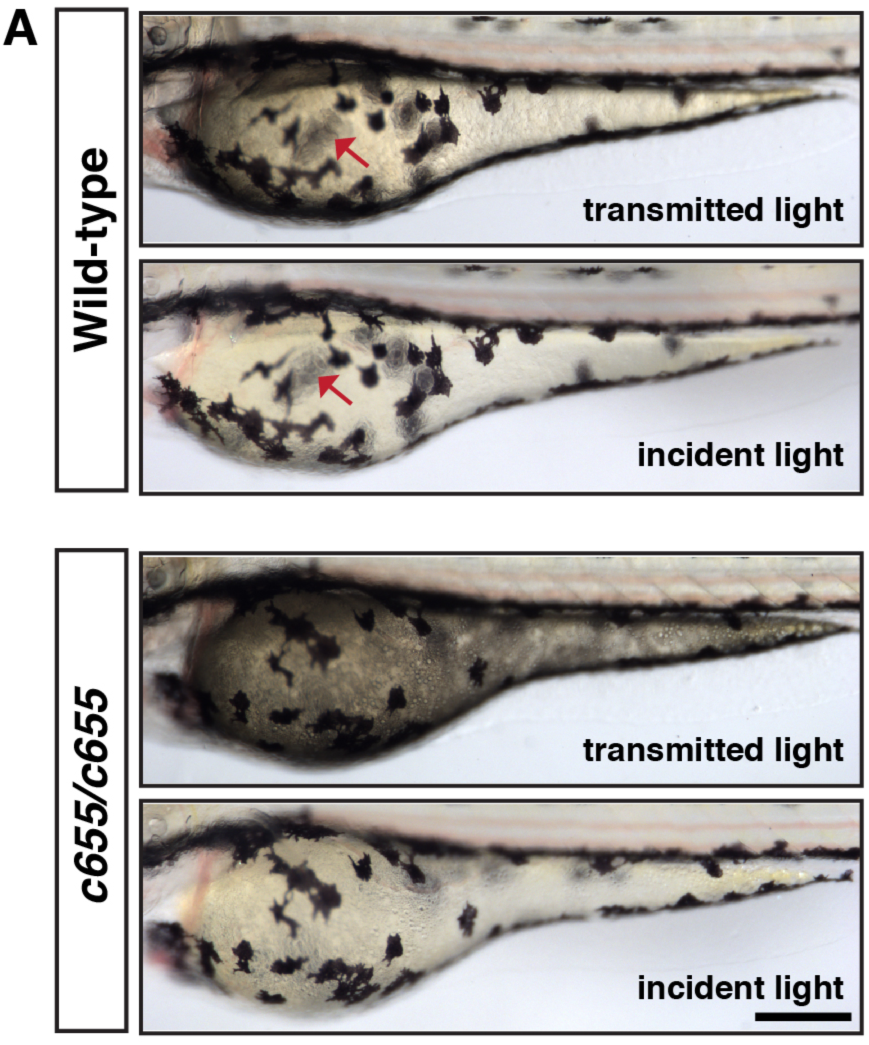
Lipid droplets block light transmission through the yolk of the embryo. (A) Wild-type and *c655* mutant embryos were imaged at 3 dpf using either transmitted light (illumination below the fish) or incident light (illumination from above the fish). The wild-type embryos are translucent; the pigment cells on the opposite side of the embryo (red arrow) are visible through the yolk with both light sources. The yolk is opaque in the mutants; it appears dark with transmitted light and white with incident light. Pigment cells on the opposite side of the embryo are barely visible in mutant embryos, regardless of light source. Scale = 200 μM.

One of the SNVs linked to the *c655* phenotype was a missense mutation predicted to be deleterious in exon 18 of the microsomal triglyceride transfer protein gene (ENSDARG00000008637, Chr1:11,421,261 GRCz10). A previously identified missense mutation in exon 11 of zebrafish *mttp*, stalactite (*stl*), also presents with an opaque yolk phenotype (Figure 1C, top) (Avraham-Davidi et al., 2012), suggesting that the *c655* opaque yolk phenotype might result from this newly identified missense mutation in *mttp*. To test this hypothesis, we performed complementation crosses between *mttp^c655/+^* heterozygous fish and *mttp^stl/+^* heterozygous fish. The *c655* mutation failed to complement the *mttp^stl^* mutation, as one-quarter of the embryos in these crosses displayed the opaque yolk phenotype (Figure 1C, bottom). All of the embryos exhibiting opaque yolks were heterozygous for both the *mttp^stl^* and the *c655* mutation in *mttp*. This strongly argues that the G/T SNV in exon 18 of *mttp* is the causative allele for the *c655* opaque yolk phenotype. This was further confirmed by rescuing the *c655* opaque yolk phenotype with injections of a wild-type mttp-FLAG plasmid at the 1-cell stage (Figure 1 – figure supplement 2).

**Figure 1 – figure supplement 2:**
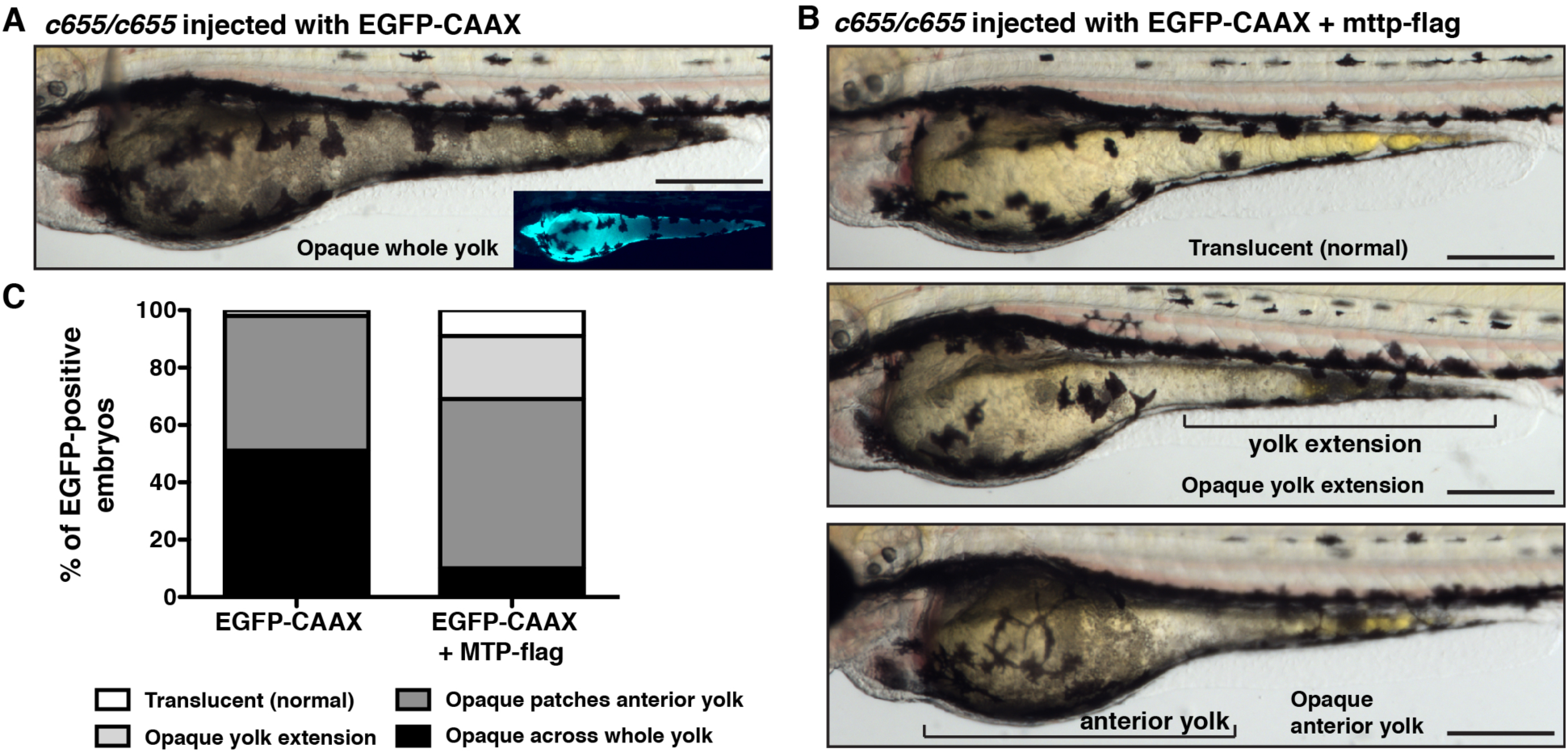
Expression of wild-type zebrafish *mttp*-FLAG rescues the opaque yolk phenotype in *mttp^c655/c655^* embryos. One-cell stage *mttp^c655/c655^* embryos were co-injected with CMV: *mttp*-FLAG and the CMV: *eGFP-CAAX* plasmid, or CMV: *eGFP-CAAX* alone as a control. Embryos expressing eGFP in the YSL were imaged at 3 dpf, and images were blinded and scored for the degree of yolk opacity. (A) Representative image of an *mttp^c655/c655^* mutant embryo expressing eGFP-CAAX in the YSL and a fully opaque yolk. (B) Examples of injected embryos with varying degrees of yolk opacity (normal translucent yolk, opaque region in the yolk extension, opaque patches in the anterior yolk with or without opaque yolk extension). (C) Images were binned into the four noted categories of yolk opacity. Results represent pooled data from 3 independent experiments, n = 91 control and 102 Mttp-FLAG eGFP-positive embryos total. Chi-square test, p<0.001. Scale = 500 μM.

Both the *mttp^stl^* allele and *mttp^c655^* allele are missense mutations. The *stl* allele results in the conversion of a leucine to a proline at residue 475 and the *c655* mutation is a glycine to valine mutation in the C-terminus of the protein at residue 863 (total length = 884 residues) (Figure 1D). An additional SNV in *mttp* at position Chr1:11,421,300 GRCz10 (T/C) causing a missense mutation (M850T) was identified in *c655* mutants; however, this SNP was not predicted to be deleterious and has been previously noted in the Ensembl zebrafish genome database. Furthermore, no change in mRNA expression was noted for *mttp* in the *mttp^c655^* mutants in our RNAseq data-set (log2[fold change] = 0.18, adj. p-value = 0.19).

Although the *mttp^stl/stl^*, *mttp^c655/c655^,* and trans-heterozygous *mttp^stl/c655^* fish all exhibit opaque yolks, the *mttp^stl/stl^* mutants have a more severe phenotype, in that their yolks are darker and they retain the opaque phenotype longer during development. The *mttp^c655/c655^* mutant phenotype is the least severe and the trans-heterozygotes exhibit an intermediate phenotype (Figure 1 – figure supplement 3). Encouraged by the differences in embryonic phenotype, we hypothesized that these mutations would provide an opportunity to further dissect the molecular details of MTP function *in vivo*.

**Figure 1 — figure supplement 3:**
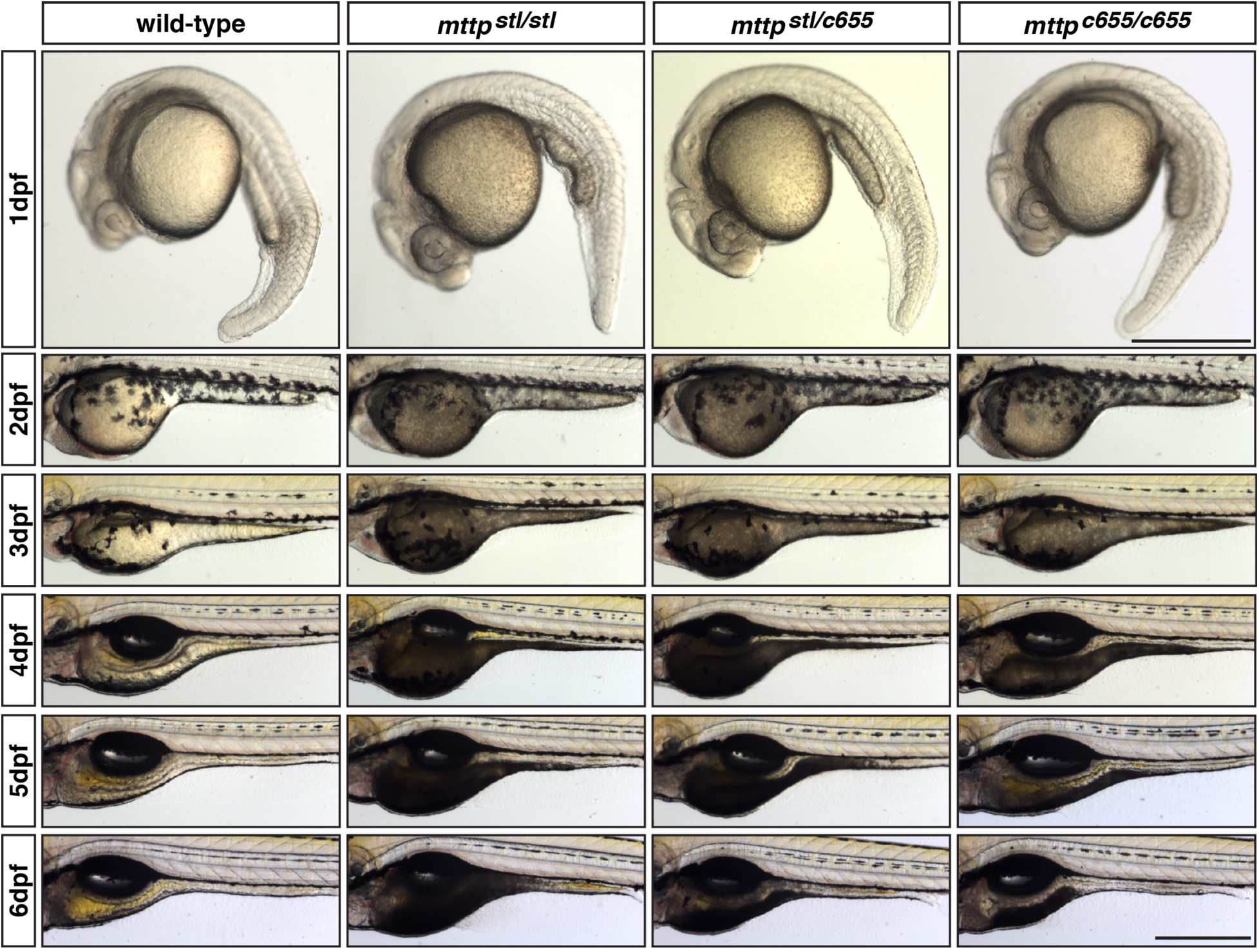
The *stl* and *c655* mttp mutations have differential effects on the degree of yolk opacity during embryonic development. Representative images of wild-type, *mttp^stl/stl^*, *mttp^c655/c655^* and trans-heterozygous *mttp ^stl/c655^* mutants from 1 dpf to 6 dpf. The *mttp^stl/stl^* mutants are visibly opaque at 1 dpf and the area of opacity is retained for longer than in *mttp^stl/c655^* or *mttp^c655/c655^* mutants. 3 dpf images are the same fish shown in Figure 1. Scale = 500 μM.

### Mttp mutants accumulate cytoplasmic lipid droplets in the yolk syncytial layer

As a lecithotrophic organism, zebrafish rely on their maternally-derived yolk as the source of nutrients and building blocks for embryogenesis (Hiramatsu et al., 2015; Mani-Ponset et al., 1996; Vernier and Sire, 1977). The yolk is rich in lipids (Fraher et al., 2016; Miyares et al., 2014; Wiegand, 1996) and following lipolysis and re-esterification, the lipids are packaged into lipoproteins in the ER of the yolk syncytial layer (YSL), a multi-nucleated cytoplasm that surrounds the yolk mass (Figure 2A)(Carvalho and Heisenberg, 2010; Kimmel and Law, 1985; Walzer and Schonenberger, 1979a, b). The zebrafish YSL expresses apolipoprotein B (Otis et al., 2015) and microsomal triglyceride transfer protein (Marza et al., 2005; Schlegel and Stainier, 2006). The YSL produces B-lps (Thierer et al., In Press; Vernier and Sire, 1977; Walzer and Schonenberger, 1979a), similar to the intestine (Glickman et al., 1976; Kessler et al., 1970), liver (Mahley et al., 1970), placenta (Madsen et al., 2004), and embryonic yolk sac in mammals (Farese et al., 1996; Plonne et al., 1992). In the mammalian intestine, ApoB mRNA is edited, resulting in a truncated ApoB molecule (ApoB48)(Davidson and Shelness, 2000; Kane et al., 1980); however, there is no evidence for editing in zebrafish, so all B-lp-producing tissues, including the YSL, secrete lipoproteins containing ApoB100 (Thierer et al., In Press).

**Figure 2:**
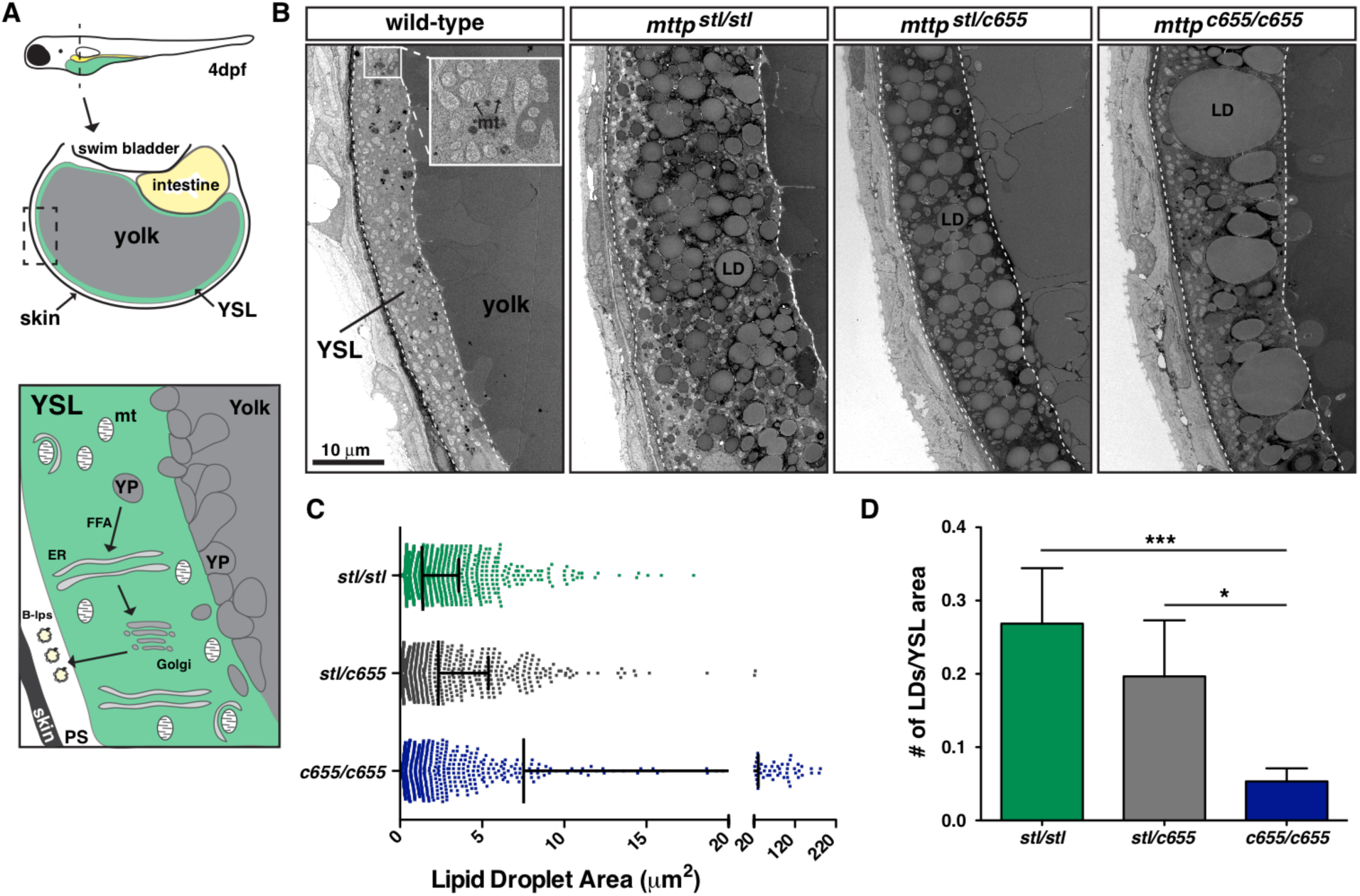
The opaque yolk phenotype results from the accumulation of aberrant cytoplasmic lipid droplets in the yolk syncytial layer. (A) (Top) Cartoon depicting the cross-sectional view of a 4-dpf zebrafish embryo. The yolk syncytial layer (YSL) surrounds the yolk mass and serves as the embryonic digestive organ. The dashed box indicates the view expanded in the bottom panel and in panel B. (Bottom) Stored yolk lipids undergo lipolysis in yolk platelets (YP), presumably releasing free fatty acids into the YSL. These fatty acids are re-esterified in the ER bilayer to form triglycerides, phospholipids, and cholesterol esters. The lipids are packaged into ApoB-containing lipoproteins in the ER with the help of MTP and are further processed in the Golgi before being secreted into the perivitelline space (PS) and then circulation. (B) Representative transmission electron micrographs of the yolk and YSL from wild-type and *mttp* mutants; dashed lines delineate the YSL region, mt = mitochondria, scale = 10 μm. (C) Quantification of lipid droplet size in *mttp* mutants, n ≥ 700 lipid droplets in 2 fish per genotype; mean +/-SD. (D) Quantification of the number of lipid droplets per YSL area, n = 7-9 YSL regions per genotype (3–5 regions per fish, 2 fish per genotype); mean +/-SD, Kruskall-Wallis with Dunn’s Multiple Comparison test, vs. *mttp^c655/c655^* * p < 0.05, *** p < 0.001.

**Figure 2 – figure supplement 1:**
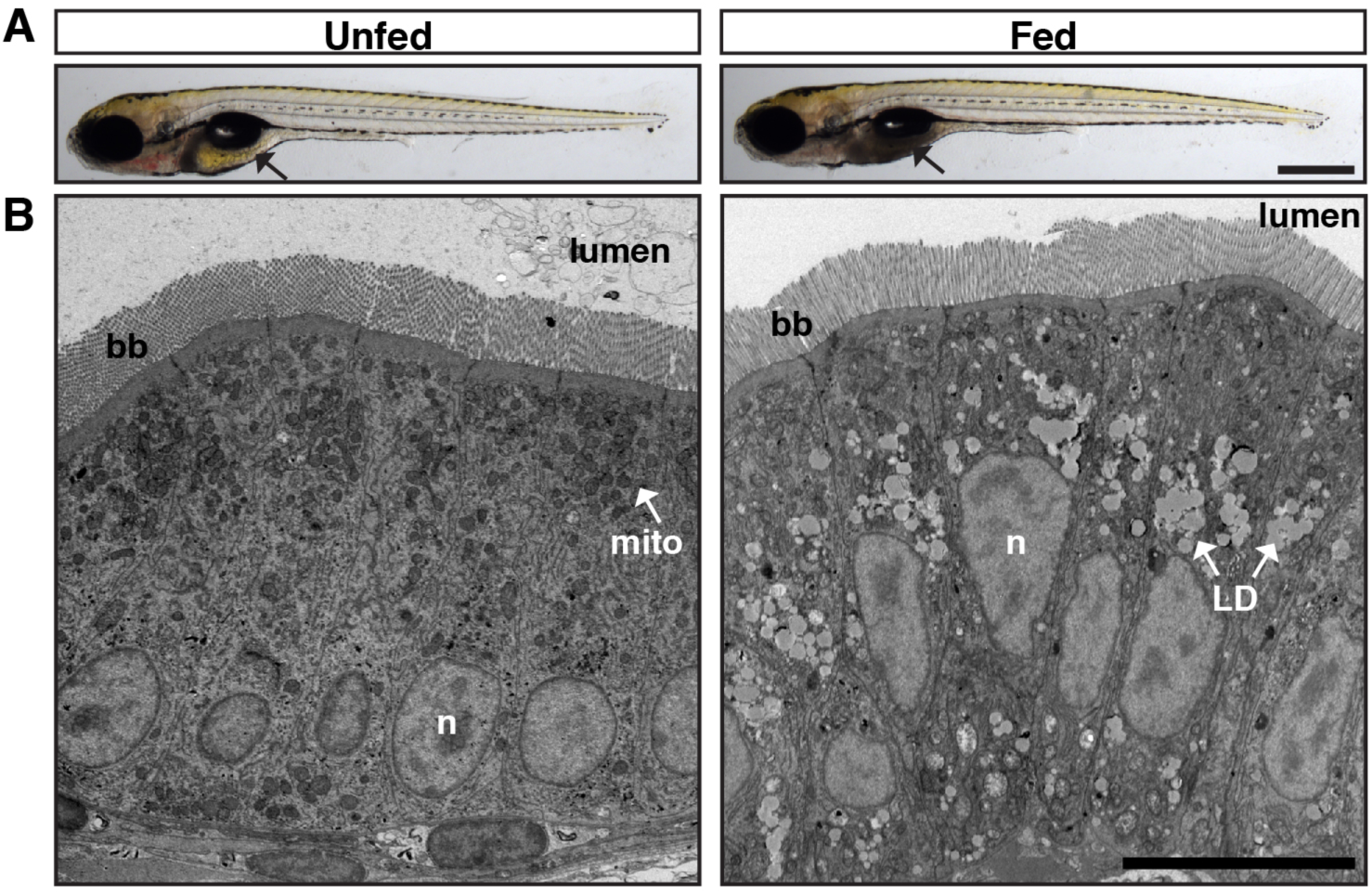
Lipid droplets block light transmission through the larval intestine. (A) Wild-type fish at 6 dpf were fed a high-fat meal for 1h as described previously (Otis and Farber, 2016). Unfed fish have translucent intestines (black arrow, left) when imaged with transmitted light, whereas fed fish have opaque intestines (black arrow, right). Scale = 500 μM. (B) Electron microscopy following a 1 h high-fat feed reveals an accumulation of cytoplasmic lipid droplets in the intestinal enterocytes. By scattering light and blocking light transmission through the intestine, the accumulation of cytoplasmic lipid droplets causes the intestine to appear opaque. Nucleus (n), mitochondria (mito), brush border (bb), lipid droplet (LD). Scale = 10 μM.

When MTP is mutated or absent, B-lp production is reduced or absent and TG accumulates in cytoplasmic lipid droplets (LDs) instead (Khatun et al., 2012; Raabe et al., 1999). We have previously shown that accumulation of LDs in intestinal enterocytes of zebrafish larvae fed a high-fat meal causes the gut to be opaque (Otis and Farber, 2016)(Figure 2 – figure supplement 1), most likely due to the lipid droplets’ ability to scatter light (Hwang et al., 2018; Michels et al., 2008). Therefore, we hypothesized that the yolk opacity in the *mttp* mutant embryos is due to aberrant accumulation of LDs in the cytoplasm of the YSL. Using transmission electron microscopy, we found that the YSL in the wild-type embryos contains very few, if any, canonical YSL LDs, whereas the *mttp^stl/stl^*, *mttp^c655/c655^* and trans-heterozygous *mttp^stl/c655^* embryos accumulate substantial numbers of cytoplasmic LDs (Figure 2B). LDs in *mttp^stl/stl^* mutants are more numerous and more uniform in size, whereas the *mttp^c655/c655^* mutants often had very large LDs in addition to small droplets (Figure 2C). As a result, the number of LDs per area of the YSL is reduced in the *mttp^c655/c655^* mutants compared to *mttp^stl/stl^* mutants (Figure 2D). The trans-heterozygous fish had LDs that were more similar in size to the *mttp^stl/stl^* mutants and had a trend toward fewer lipid droplets per YSL area, although this was not significant.

### The stl and c655 mutations have differential effects on adult size and steatosis

Patients with Abetalipoproteinemia often present in infancy with growth retardation, diarrhea, fat malabsorption, and failure to thrive (reviewed in (Lee and Hegele, 2014)), and whole body deficiency of MTP in a murine model is embryonic lethal (Raabe et al., 1998). In agreement with the mouse phenotype, in the original characterization of the zebrafish *mttp^stl/stl^* phenotype it was noted that the fish did not survive past 6 days post fertilization (dpf)(Avraham-Davidi et al., 2012). Therefore, we were surprised to find that some of the *stl* mutants were able to survive past larval stages. While these fish are generally much smaller in length and mass (Figure 3A,B) and their viability is reduced relative to their siblings (expected 25%, observed 3.8% [5/131 fish] at 7.5 months), some of these fish have survived to be at least 18 months old. Survival rates are better when the mutants are reared separately and are not competing with siblings for food. Although these fish can reproduce, this is rare. In contrast, the *mttp^c655/c655^* mutants do not exhibit reduced viability and we did not find any reduction in size or fertility of the *c655* mutants compared with siblings (Figure 3A,B). No difference in length or mass was noted in fish trans-heterozygous for *mttp^stl/c655^* (Figure 3A,B).

**Figure 3:**
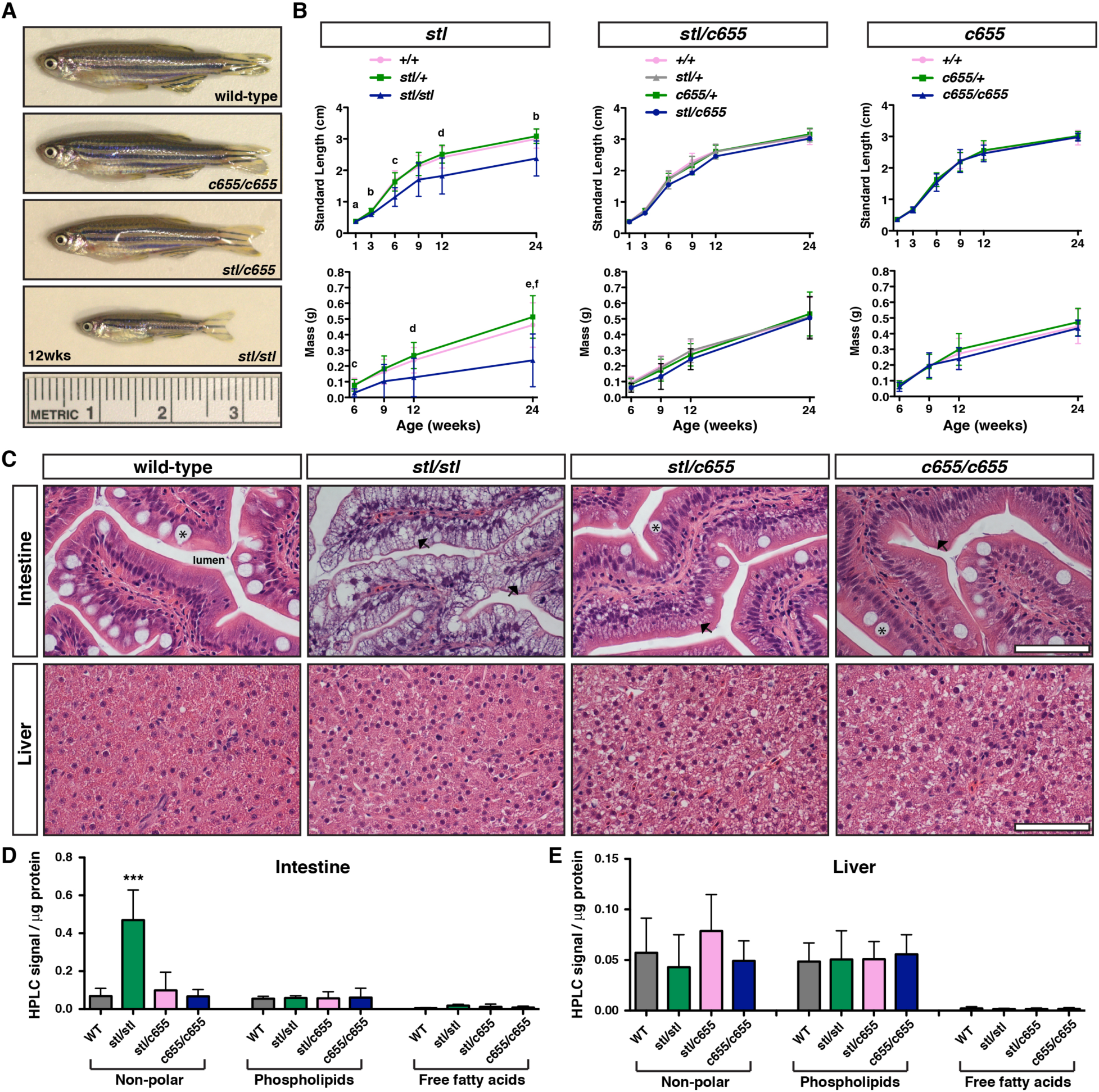
The *stl* and *c655 mttp* mutations have differential effects on the growth and accumulation of lipid in intestine. (A) Representative images of WT and *mttp* mutant fish at 12 weeks of age. (B) Developmental time-course of standard length and mass measurements of *mttp* mutant fish and siblings. Results are representative of pooled data from two independent experiments, n = 7–80 fish/genotype/time-point, mean +/-SD. Significance was determined with a Robust ANOVA and Games-Howell post-hoc tests were used to make pair-wise comparisons at each time point. Using a Bonferroni correction, p-values were adjusted to control for multiple comparisons (6 length or 4 mass comparisons), a: *stl/+* vs. *stl/stl*, p <0.01; b: *+/+* vs. *stl/stl* and *stl/+* vs. *stl/stl*, p <0.05; c: *+/+* vs. *stl/stl* and *stl/+* vs. *stl/stl*, p <0.001; d: *stl/+* vs. *stl/stl*, p <0.05; e: *+/+* vs. *stl/stl*, p <0.01; f: *stl/+* vs. *stl/stl*, p<0.001. (C) Representative images of H&E stained intestine and liver from adult WT and *mttp* mutant fish (7.5 months), scale = 50 μm, * indicate goblet cell, arrows indicate representative lipid accumulation in enterocytes. (D & E) Quantification of non-polar lipids, phospholipids, and free fatty acids in intestine and liver tissue using high-performance liquid chromatography; n = 5–6 fish per genotype, mean +/-SD, One-way ANOVA with Bonferroni post-hoc tests were performed separately for each lipid class; *** p < 0.001 *stl/stl* vs. all other genotypes.

**Figure 3 – figure supplement 1:**
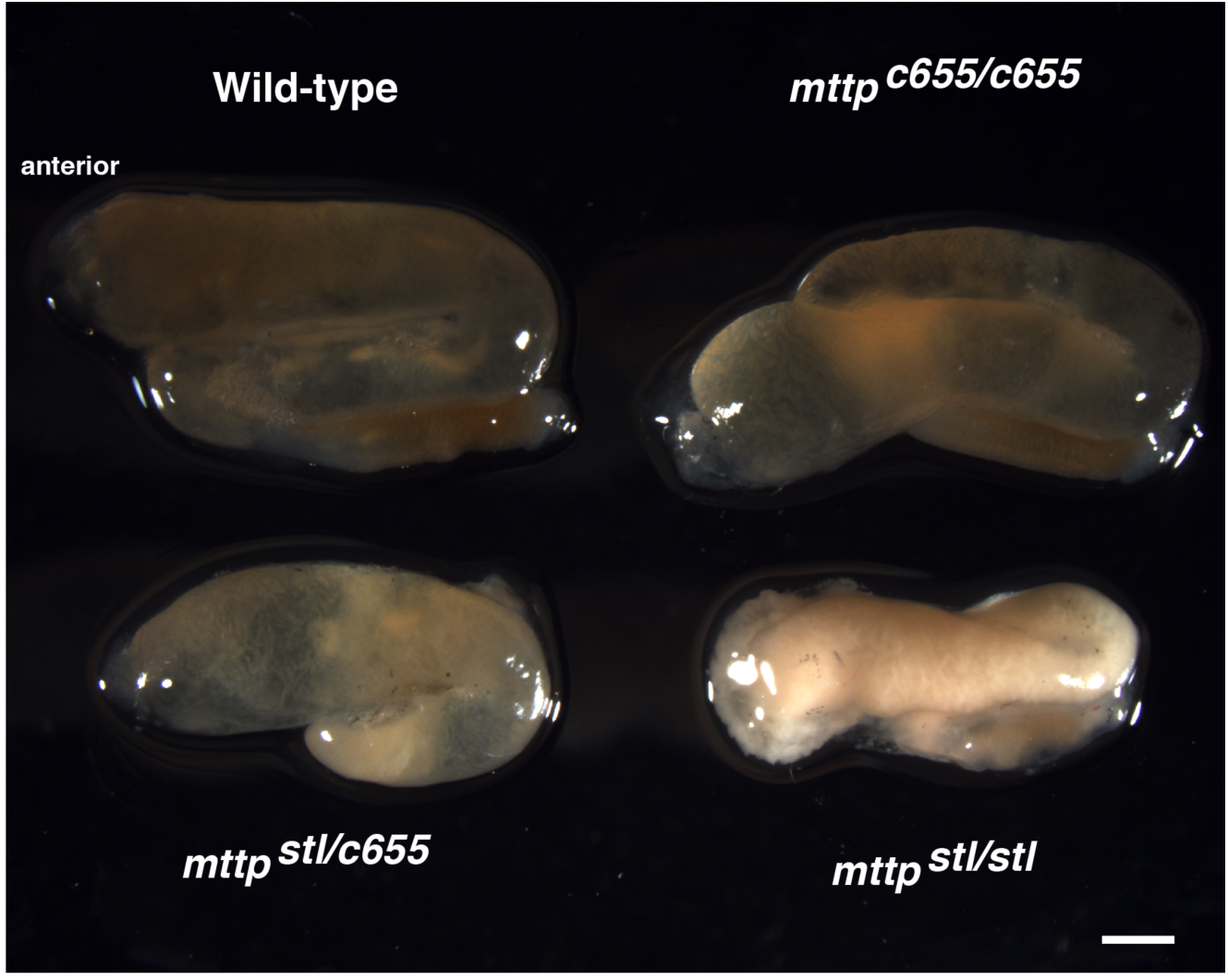
Significant lipid accumulation in the intestine of *stl/stl* but not in *c655/c655* mutants. Representative images of isolated intestines from adult WT and *mttp* mutant fish (7.5 months), scale = 1 mm.

To assess why the *mttp^stl/stl^* mutants, but not the *mttp^c655/c655^* mutants, have defects in growth, we examined whether the mutations have differential effects on fat malabsorption. H&E staining of intestinal tissue from fasted adults revealed gross accumulation of lipid in the cytoplasm of enterocytes in the *mttp^stl/stl^* fish (Figure 3C, Figure 3 – figure supplement 1). The *mttp^c655/c655^* mutants were largely protected from this abnormal lipid retention. While the trans-heterozygous fish exhibited more visible lipid retention than either wild-type or *mttp^c655/c655^* fish, this was not as profound as in the *mttp^stl/stl^* fish. Retention of non-polar lipid (TG and cholesterol ester), phospholipid, and free fatty acids in the intestine were also quantified using HPLC and the *mttp^stl/stl^* mutants contain ∼7 times more neutral lipid than *c655* mutants (Figure 3D). This data suggests the growth defects in *mttp^stl/stl^* mutants result from defects in dietary lipid absorption in the intestine.

Besides accumulating lipids in the intestine, Abetaliporoteinemia patients sometimes develop hepatic steatosis (reviewed in (Lee and Hegele, 2014)). Similarly, hepatocyte-specific deficiency of MTTP in mice causes TG and cholesterol to accumulate in the liver (Khatun et al., 2012; Raabe et al., 1999). Therefore, we hypothesized that the adult zebrafish *mttp* mutants would also exhibit liver steatosis. However, mutants were not different than wild-type (Figure 3C,D). Moreover, there was no significant difference in measured lipids in the livers of the different fish (Figure 3E). While we were surprised that the *mttp^stl/stl^* mutants had very little accumulation of lipid in their livers, this is in agreement with findings that combined intestinal and liver deficiency of Mttp in mice results in accumulation of TG in the intestine, but not in the liver (Iqbal et al., 2015).

### Mttp mutations reduce the size and number of ApoB-containing lipoproteins in vivo

To understand how the *mttp* mutations affect the production and size of B-lps during embryonic development, we crossed the *mttp^stl^* and *mttp^c655^* mutations into our LipoGlo reporter line (Thierer et al., In Press). These fish express an in-frame fusion of the engineered luciferase reporter Nanoluc at the C-terminus of the Apolipoprotein Bb.1 gene (Figure 4A). Since ApoB is an obligate structural component of B-lps with only one copy per lipoprotein particle (Elovson et al., 1988), the relative number of tagged lipoprotein particles can be quantified using the Nano-Glo assay in extracts from transgenic fish (Thierer et al., In Press). B-lp levels were measured in whole fish lysate throughout embryonic development from 2–6 dpf. During this time, the fish are relying solely on yolk lipids and the ApoB quantity measurements reflect lipoprotein particles in the secretory pathway in the YSL, in the circulation, and in cells prior to degradation of endocytosed lipoproteins. Wild-type embryos exhibit an increase in B-lp particle number from 2–3 dpf as yolk lipid is packaged into lipoproteins, and then numbers decline as the yolk is depleted and the lipids in the lipoproteins are taken up by target tissues and lipoprotein particles are degraded (Figure 4B). The *mttp^c655/c655^* embryos have the same relative number of ApoB particles as wild-type embryos at 2 dpf, but from 3-6 dpf the numbers of particles never reach wild-type levels and decline more rapidly. In contrast, *mttp^stl/stl^* embryos have profound defects in B-lp production, since amount of ApoB is significantly lower than wild-type siblings at 2 dpf (Figure 4B) (Thierer et al., In Press).

**Figure 4:**
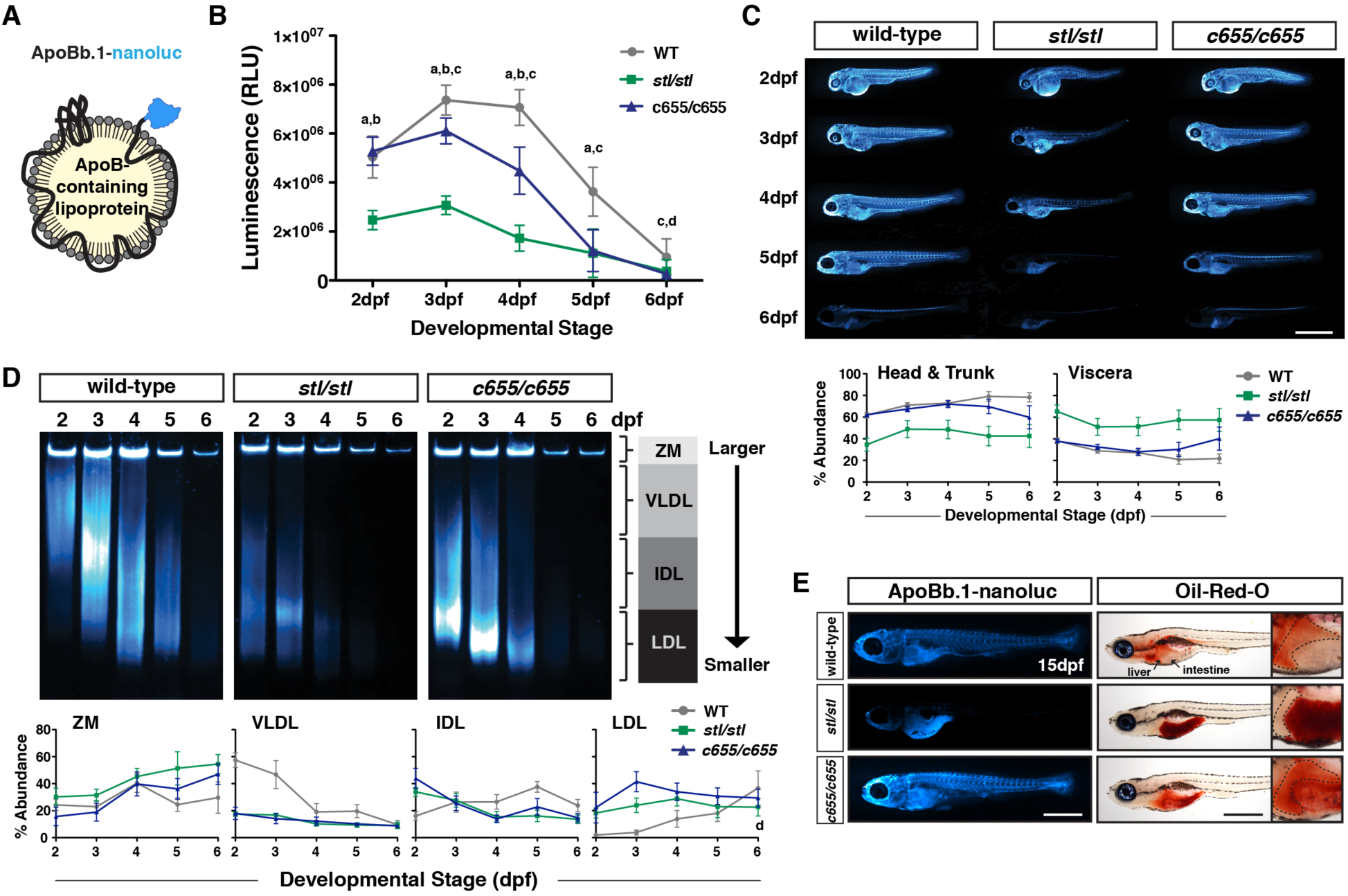
The *stl* and *c655* mttp mutations have differential effects on ApoB-containing lipoprotein number, size and distribution *in vivo*. (A) LipoGlo fish express the Nanoluc® luciferase enzyme as a C-terminal fusion on ApoBb.1 as a result of TALEN-based genomic engineering. (B) LipoGlo signal (RLU: relative luminescence units) in WT, *mttp^stl/stl^* and *mttp^c655/c655^* fish throughout embryonic development (2–6 dpf). Results represent pooled data from 3 independent experiments, n = 22–34 fish/genotype/time-point. Significance was determined with a Robust ANOVA, Games-Howell post-hoc tests were performed to compare genotypes at each day of development, and p-values were adjusted to control for multiple comparisons, a = WT vs. *mttp^stl/stl^*, p<0.001, b = *mttp^c655/c655^* vs. *mttp^stl/stl^*, p<0.001, c = WT vs. *mttp^c655/c655^*, p<0.001, d = WT vs. *mttp^stl/stl^*, p<0.05. (C) Representative whole-mount images of B-lp localization using LipoGlo chemiluminescent microscopy in WT, *mttp^stl/stl^,* and *mttp^c655/c655^* fish throughout development; scale = 1 mm. Graphs represent pooled data from 3 independent experiments, n = 13–19 fish/genotype/time-point; *mttp^stl/stl^* had a significantly different ApoB localization from WT and *c655/c655*, p<0.001, Robust ANOVA. Post-hoc analysis reveals statistical differences at all developmental stages p<0.05–0.001 (Games-Howell). (D) Representative LipoGlo PAGE gels and quantification of B-lp size distribution from whole embryo lysates during development. B-lps are divided into four classes based on mobility, including zero mobility (ZM), and three classes of serum B-lps (VLDL, IDL and LDL). Graphs show subclass abundance for WT, *mttp^stl/stl^*, and *mttp^c655/c655^* fish at each day of embryonic development as described in (Thierer et al., In Press). Results represent pooled data from n = 9 samples/genotype/time-point; at each particle class size, there were statistically significant differences between genotypes (Robust ANOVA, p<0.001). Games-Howell post-hoc analysis revealed numerous differences between genotypes at each developmental stage, see figure supplement 2. (E) Representative whole-mount images of LipoGlo microscopy and Oil Red O imaging in 15-dpf embryos chow-fed for 10 days and fasted ∼18hrs prior to fixation; scale = 1 mm. Livers (outlined) are magnified for clarity in insets on right. Results represent pooled data from 3 independent experiments, n = 15 fish/genotype/time-point.

To assess the localization of the B-lps throughout the embryos during development, we fixed the embryos expressing ApoBb.1-nanoluc and performed chemiluminescent whole-mount imaging (Figure 4C). Wild-type and *mttp^c655/c655^* embryos exhibit a similar distribution pattern of LipoGlo throughout 2–4 dpf, but consistent with the quantitative assay, the signal in the head and trunk decline more rapidly in *mttp^c655/c655^* fish. By 6 dpf, both wild-type and *mttp^c655/c655^* fish show an accumulation of ApoB in the liver and the spinal cord (Figure 4C) (Thierer et al., In Press). The LipoGlo in the *mttp^stl/stl^* embryos shows a very different pattern, in that it is predominantly localized to the YSL/viscera at all stages and is at very low levels throughout the rest of the body (Figure 4C). This is consistent with the prolonged retention of the opaque yolk phenotype (Figure 1 – figure supplement 3) and suggests that the *mttp^stl/stl^* mutants are more defective at secreting B-lps from the yolk than the *mttp^c655/c655^* mutants.

To determine whether the *mttp* mutations alter the size distribution of B-lps, we performed native polyacrylamide gel electrophoresis of larval homogenates expressing the LipoGlo reporter. Following electrophoretic separation and chemiluminescent imaging of the gels, B-lps were classified into four different classes based on their migration distance (zero mobility, very low-density lipoproteins (VLDL), intermediate-density lipoproteins (IDL), or low-density lipoproteins (LDL)(Thierer et al., In Press). During development, the pattern of B-lps in wild-type embryos is initially defined by VLDL (2 dpf), but expands to include IDL and LDL by 3–4 dpf as the VLDL particles produced by the YSL are lipolyzed by the body tissues (Figure 4D, Figure 4 — figure supplement 1, (Thierer et al., In Press)). By 5–6 dpf, the yolk is depleted and any remaining small particles are degraded. In contrast, both *mttp^stl/stl^* and *mttp^c655/c655^* embryos produce very few VLDL particles (Figure 4D, 2 dpf), and instead, produce predominantly IDL and LDL-sized particles.

**Figure 4 – figure supplement 1:**
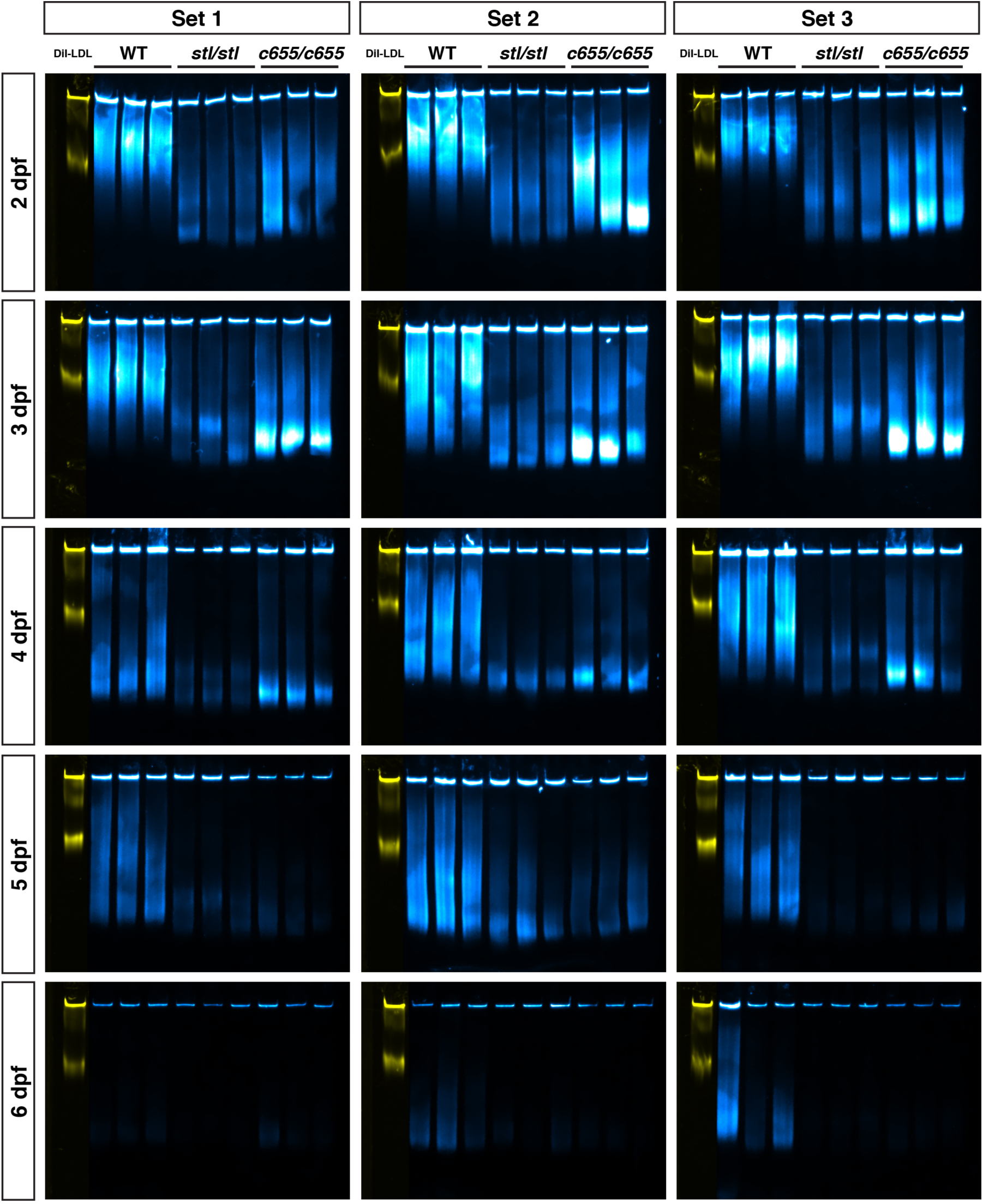
LipoGlo lipoprotein gel primary data. Original gels corresponding to the data in Figure 4D. Each gel shows a composite image of the fluorescent DiI-LDL migration standard (yellow) and LipoGlo emission chemiluminescent exposure (blue) from WT, *mttp^stl/stl^* and *mttp^c655/c655^* fish. Gels were analyzed as detailed in (Thierer et al., In Press) and lipoprotein particles were binned into four classes based on migration relative to the DiI-LDL standard, including zero mobility (ZM), and three classes of serum B-lps (VLDL, IDL and LDL).

**Figure 4 – figure supplement 2:**
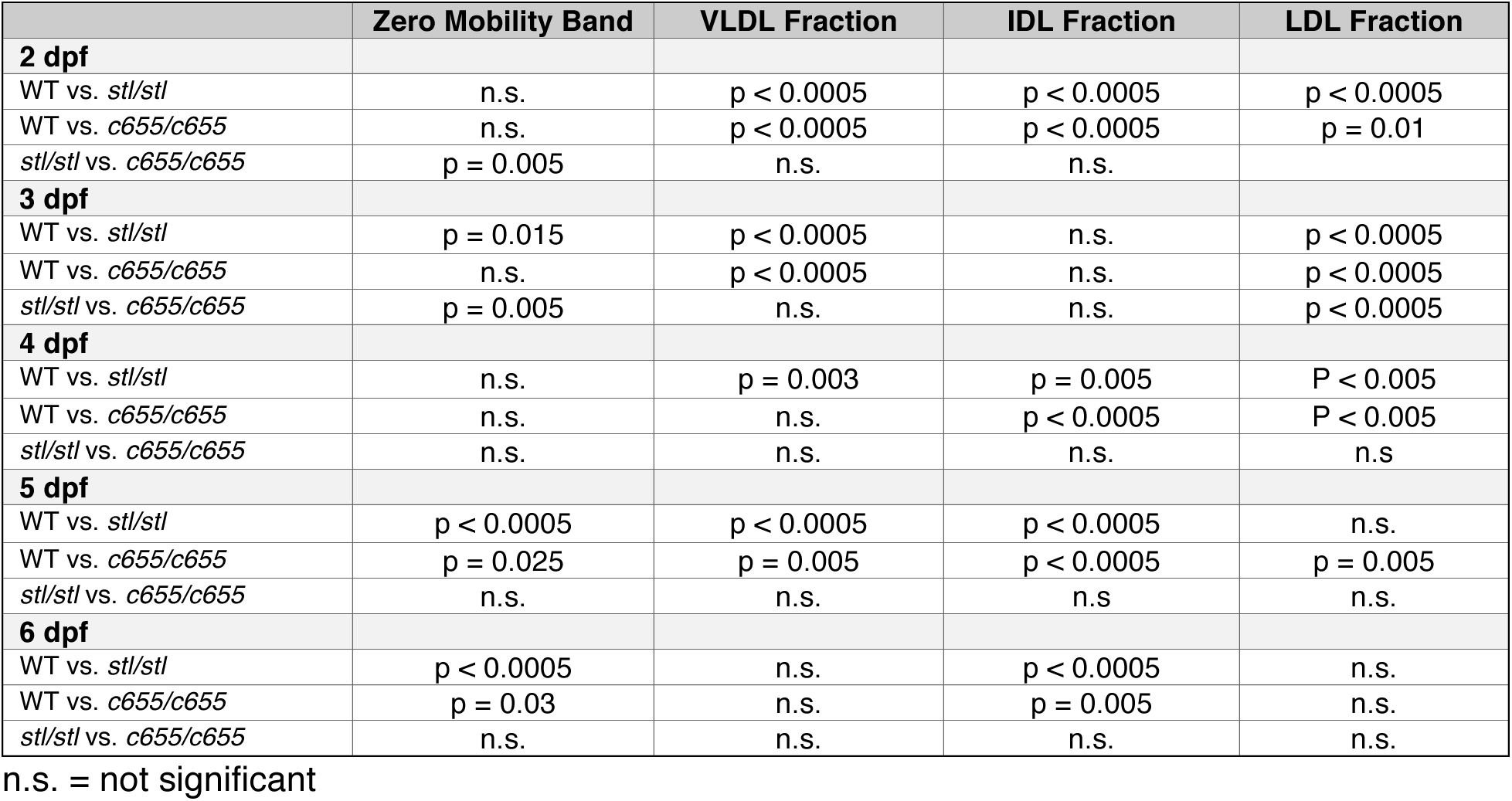

Taken together, the LipoGlo data from embryos indicates that the *mttp^stl/stl^* and *mttp^c655/c655^* embryos are producing and secreting fewer, smaller B-lps from the yolk, although the effect is more severe in *stl* mutants. Given the gross accumulation of lipid in the intestines of the adult *mttp^stl/stl^* mutants, compared to the *mttp^c655/c655^* fish (Figure 3C), we hypothesize that the *mttp^stl/stl^* mutants are also less effective at secreting chylomicrons from the enterocytes. To test this hypothesis, we performed chemiluminescent imaging using the LipoGlo reporter in 15-dpf larvae fed a chow diet for 10 days and then fasted overnight. Wild-type LipoGlo fish have abundant ApoB throughout their circulation and tissues (73.1+/-4.0% in head and trunk vs. 26.9 +/-4.0% in viscera, mean +/-SD, n = 15 fish) (Figure 4E). The *mttp^c655/c655^* mutation does not prevent secretion of ApoB to the body tissues (73.1 +/-3.7% in head and trunk vs. 26.9 +/-3.7% in viscera). In contrast, the *mttp^stl/stl^* fish have abundant LipoGlo signal in their intestine, and much less in other tissues compared to WT (41% +/-11% in head and trunk vs. 59 +/-11% in viscera, p<0.001, Kruskall-Wallis & Dunn’s Multiple Comparisons Test) (Figure 4E). In agreement, staining the neutral lipid with Oil Red O indicates *mttp^stl/stl^* mutants retain substantial lipid in their intestines, whereas *mttp^c655/c655^* mutant fish have less lipid remaining in their intestine, but do accumulate some lipid in their livers (Figure 4E). This data argues that the *stl* mutation severely reduces B-lp secretion, not only from the yolk, but also from the enterocytes, whereas the *c655* mutation only mildly decreases ApoB secretion in both embryos and larvae.

### The c655 mutation in zebrafish mttp disrupts the triglyceride transfer activity but not the phospholipid transfer activity of the MTP complex.

The dissimilar phenotypes of *in vivo* B-lp secretion between the *stl* and *c655* mutations suggest that the two mutations are differentially affecting MTP function. To provide molecular explanations for the phenotypes observed in the different zebrafish mutants and to dissect further how each of the mutations affects MTP function, we used cell and *in-vitro*-based assays of MTP function. COS-7 cells expressing human ApoB48 were co-transfected with either an empty vector (pcDNA3), or a vector containing wild-type zebrafish *mttp*, *mttp-stl* or *mttp-c655*, all with C-terminal FLAG-tags. We found that *stl* and *c655* expressing COS-7 cells have significantly reduced concentrations of ApoB in the conditioned media compared to wild-type Mttp expressing cells (Figure 5A). ApoB levels in the media of the *stl*-expressing cells were similar to cells transfected with empty vector, indicating background secretion levels. The concentration of ApoB48 was significantly higher in the media of *c655*-expressing COS-7 cells compared to the *stl*-expressing cells and cells transfected with empty vector (Figure 5A). ApoB48 concentrations in the cell lysate of cells expressing *stl* were significantly higher than wild-type Mttp and *c655*-expressing cells (Figure 5B). These data suggest that the *stl* mutation does not support ApoB48 secretion whereas the *c655* mutation does support ApoB48 secretion, but with reduced efficacy compared to wild-type Mttp.

**Figure 5:**
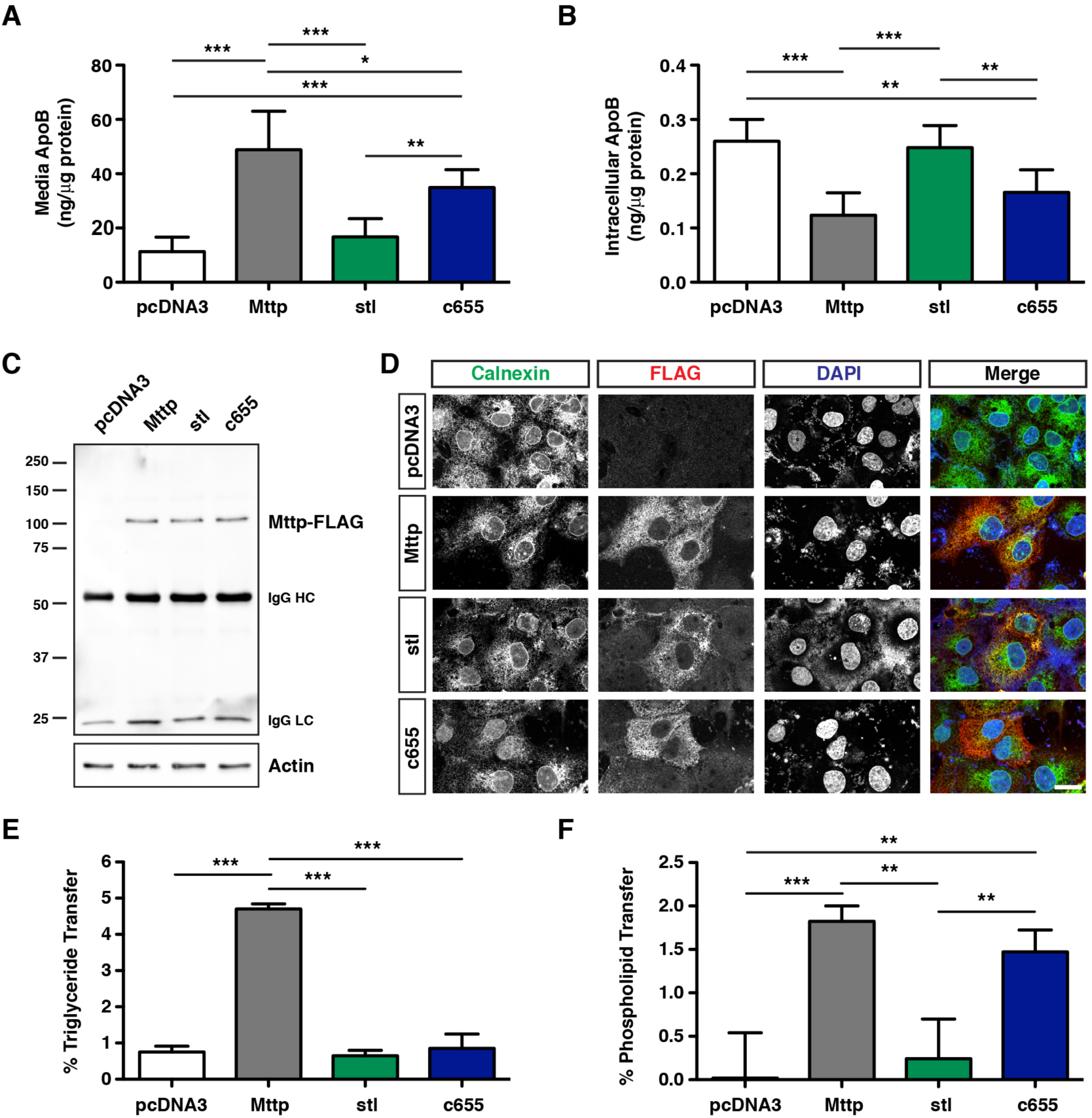
The *c655* mutation disrupts the triglyceride transfer activity, but not the phospholipid transfer activity of the zebrafish MTP complex. (A-B) COS-7 cells were first transfected with an expression vector for human apoB48 (5 μg), distributed equally in 6-well plates, and subsequently transfected with plasmids expressing either wild-type zebrafish *mttp*-FLAG, *mttp^stl^*-FLAG, *mttp^c655^*-FLAG, or empty vector (pcDNA3) (3 μg). After 72 h, ApoB48 was measured via ELISA in media (A) or in the cell (B). Data are representative of 7 independent experiments (each data point is the mean of 3 technical replicates), mean +/-SD, One-Way ANOVA with Bonferroni post-hoc tests, * p<0.05, ** p<0.01, *** p<0.001. (C) Cos-7 cells were transfected as described above and FLAG-tagged proteins were immunoprecipitated from cell lysates using anti-FLAG antibodies and eluted with FLAG peptides. Representative Western blot on eluted fractions indicates equal concentrations of the various Mttp-FLAG proteins; actin blot indicates equal loading of cell lysate. (D) Representative immunofluorescent staining using anti-FLAG (red) and anti-Calnexin (green) antibodies in COS-7 cells expressing wild-type or mutated *mttp*-FLAG constructs; scale = 25 μm. (E) COS-7 cells were transfected with plasmids expressing pcDNA3, wild-type *mttp-FLAG*, or mutant *mttp*-FLAG constructs. Cells were lysed and 60 μg of protein was used to measure the % triglyceride transfer of NBD-triolein from donor to acceptor vesicles after 45 minutes; n = 3 (each n is the mean of three technical replicates from independent experiments), mean +/-SD, One-way ANOVA with Bonferroni post-hoc tests, *** p<0.001. (F) Wild-type and mutant Mttp proteins were purified using anti-FLAG antibodies and used to measure the % transfer of NBD-labeled phosphoethanolamine triethylammonium from donor to acceptor vesicles after 180 minutes; n = 3 (each n is the mean of 3 technical replicates from independent experiments), mean +/-SD, randomized block ANOVA with Bonferroni post-hoc tests, ***p<0.001.

To eliminate the possibility that the *stl* mutation did not support ApoB48 secretion due to low expression, Mttp, *stl*, and *c655* were precipitated using anti-FLAG antibodies from cell lysates and were subjected to Western blot analysis. We found that there was no difference in the expression of wild-type Mttp and Mttp mutants in COS-7 cells (Figure 5C). Another reason for the *stl* mutation to be deficient in supporting ApoB48 secretion could be due to protein mis-localization. To check this, we immunostained Mttp-expressing COS-7 cells with anti-Calnexin and anti-FLAG antibodies. Confocal imaging shows that the wild-type Mttp and both mutant proteins were properly localized in the endoplasmic reticulum (Figure 5D). Additionally, the percentage of cells expressing the FLAG-tagged proteins were similar among all groups (Mttp-FLAG 37%, Mttp-*stl* 31%, Mttp-*c655* 41% transfection efficiency). These studies suggest that *stl* and *c655* mutant Mttp proteins are expressed to similar levels and are in the ER, where lipoprotein assembly occurs.

Next, we hypothesized that the *stl* mutant protein may not support ApoB48 secretion because it might be defective in lipid transfer activity. Triglyceride transfer assays performed using cell lysates showed that both of the mutant forms of Mttp (*stl* and *c655*) have significantly decreased triglyceride transfer activity compared to wild-type zebrafish Mttp (Figure 5E, Figure 5 – figure supplement 1a). The *stl* mutant form was also found to be defective in phospholipid transfer activity compared to wild-type (Figure 5F, Figure 5 – figure supplement 1b). Contrary to this, the *c655* mutant form had similar phospholipid transfer activity to wild-type Mttp (Figure 5F, Figure 5 – figure supplement 1b). These data suggest that the *c655* mutation in *mttp* impairs triglyceride transfer, but not phospholipid transfer activity. In contrast, the *stl* mutation is defective in both transfer activities. These differences in lipid transfer activities between the *stl* and *c655* mutations provide a biochemical explanation for their differential abilities to support ApoB48 secretion and the different phenotypes observed in the fish.

**Figure 5 – figure supplement 1:**
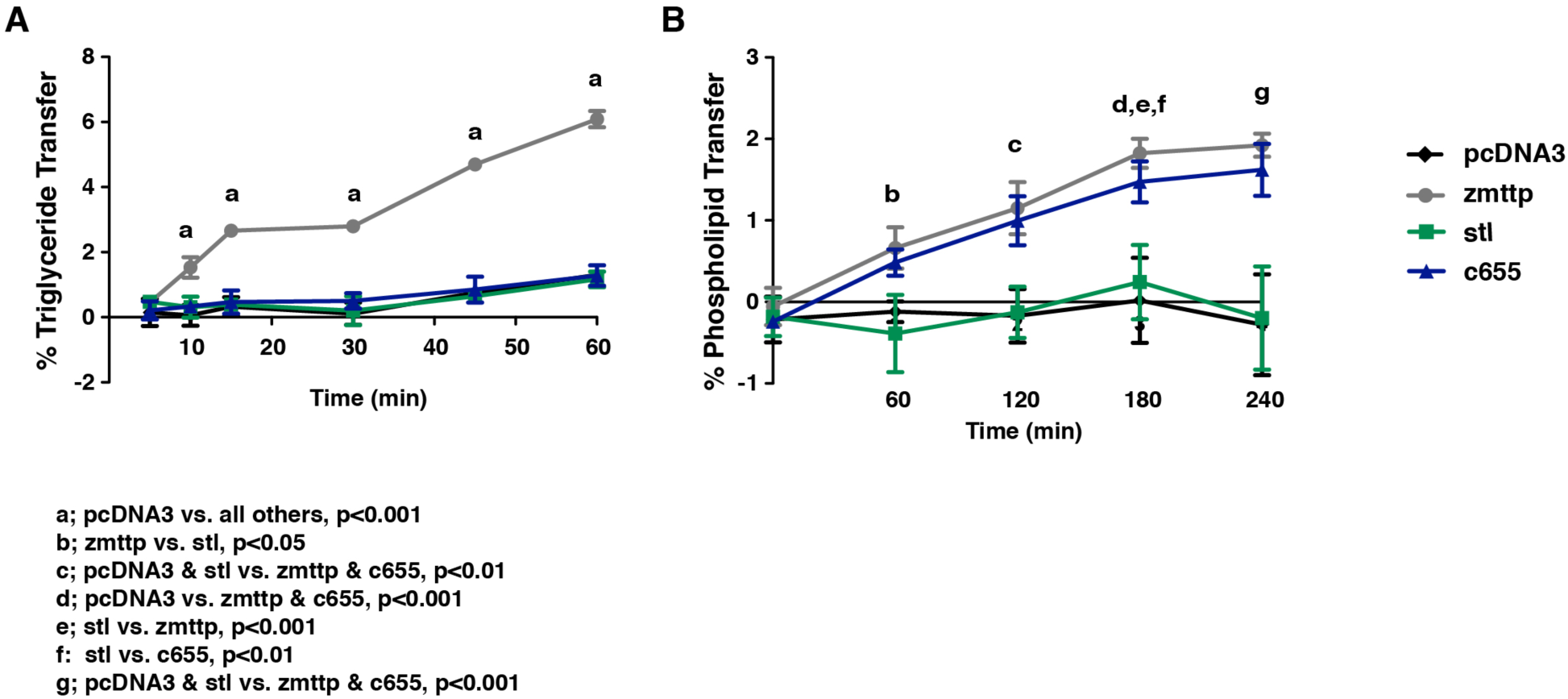
Triglyceride and phospholipid transfer assay time-course data. Measurements for triglyceride (A) and phospholipid transfer (B) by zebrafish *mttp*-FLAG and mutants over a time-course. The single time-points depicted in the bar graphs of figures 5E & F, correspond to the 45 min & 180 min (triglyceride and phospholipid transfer, respectively) time-points in the curves shown. For both, n = 3 (each n is the mean of 3 technical replicates from independent experiments), mean +/-SD, Repeated Measures ANOVA with Bonferroni post-hoc tests, significance as noted in figure.

### An orthologous c655 mutation in human MTTP (G865V) also disrupts the triglyceride transfer activity but not phospholipid transfer activity

The M subunit of MTP shares homology with lipovitellin, a lipid transfer protein in the yolk of oviparous animals (Banaszak et al., 1991). Homology modeling with the crystal structure of lamprey lipovitellin (PDB ID: 1LSH)(Anderson et al., 1998; Raag et al., 1988; Thompson and Banaszak, 2002) predicts three major structural domains in human MTTP: an N-terminal beta-barrel, a middle alpha-helical domain, and a C-terminal domain composed of two beta-sheets that form a hydrophobic lipid-binding cavity (Hussain et al., 2003a; Mann et al., 1999; Read et al., 2000). The amino acid sequence of the zebrafish Mttp is 54% identical (72% similar) to that of human MTTP and the predicted secondary and tertiary structures are highly conserved (Figure 6A).

**Figure 6:**
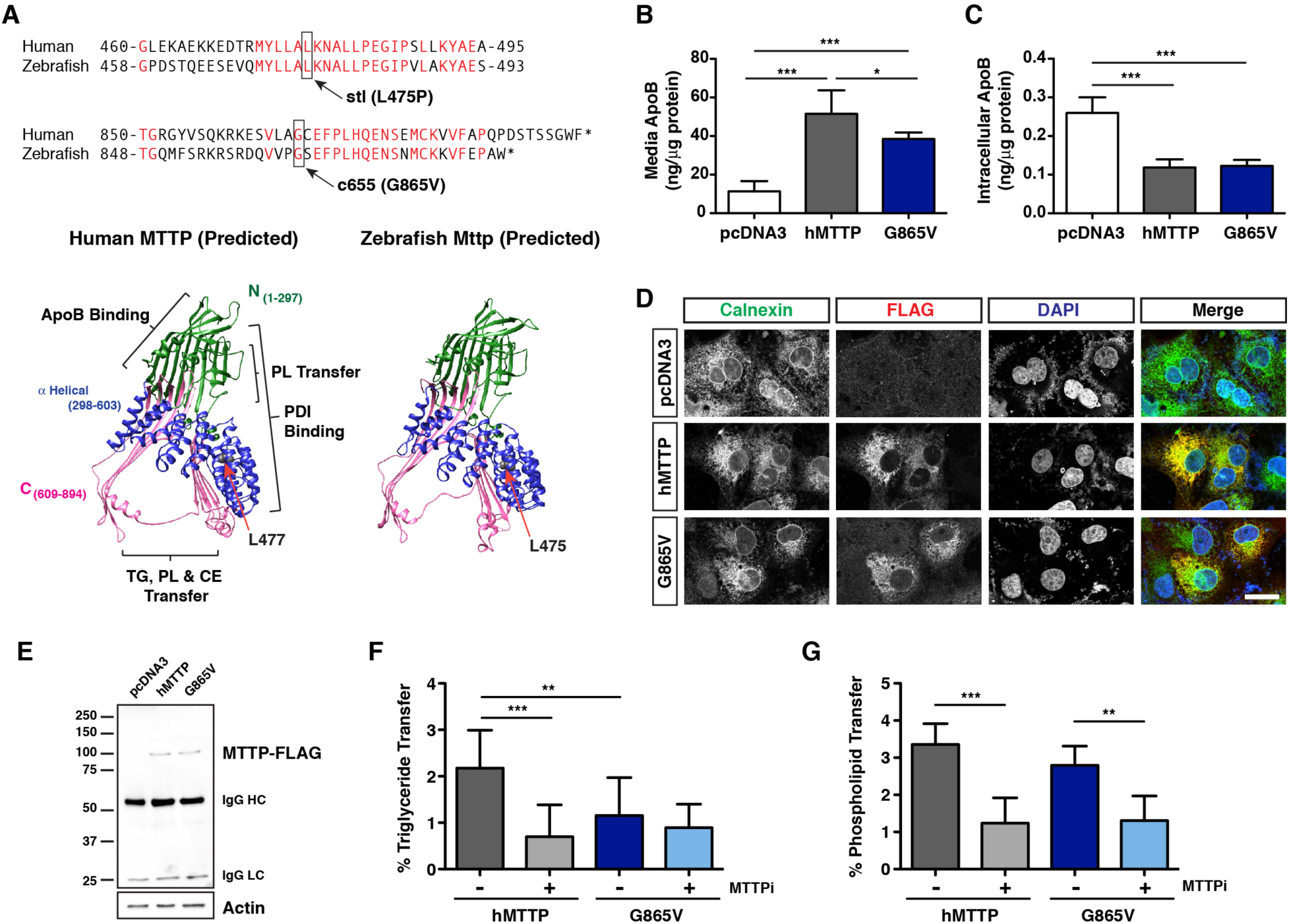
The *c655* mutation in human MTTP also disrupts the triglyceride transfer activity, but not the phospholipid transfer activity of the MTP complex. (A) Alignment of human and zebrafish Mttp amino acid sequences surrounding the *stl* or *c655* mutations. Ribbon diagrams of the predicted tertiary structures of human MTTP and zebrafish Mttp modeled using Phyre2 (Kelley et al., 2015) based on the lamprey lipovitellin structure (Anderson et al., 1998). The predicted phospholipid transfer sites in the N-terminal region (green) and the C-terminal region (pink, composed of two beta-sheets that form a hydrophobic lipid-binding pocket) are labeled. The location of the conserved human residue (L477) corresponding to the zebrafish *stl* (L475P) mutation is noted. The C-terminal region containing the *c655* mutation diverges from the lipovitellin structure and is not reliably modeled. (B,C) The *c655* mutation (G865V) was introduced into the human *MTTP*-FLAG plasmid. COS-7 cells were co-transfected with human ApoB48 and either wild-type human *MTTP*-FLAG, *MTTP*(G865V)-FLAG or empty pcDNA3 plasmids. After 72 h, apoB48 was measured via ELISA in media (B) or in the cell (C). Data are representative of 7 independent experiments (each data point is the mean of 3 technical replicates), pcDNA3 control data is re-graphed from figure 5A & 5B (data for 5A, 5B, 6B & 6C were generated together); mean +/-SD, One-Way ANOVA with Bonferroni post-hoc tests, * p<0.05, *** p<0.001. (D) Immunofluoresence in COS-7 cells expressing wild-type human MTTP-FLAG or human MTTP(G865V)-FLAG using anti-FLAG (red) and anti-Calnexin (green) antibodies; scale = 25 μm. (E) COS-7 cells were transfected and FLAG-tagged human MTTP proteins were immunoprecipitated from cell lysates using anti-FLAG antibodies and eluted with FLAG peptides. Representative Western blot on eluted fractions indicates equal concentrations of the various MTTP-FLAG proteins; actin blot indicates equal loading of cell lysate. (F) COS-7 cells were transfected with plasmids expressing human wild-type or *MTTP*(G865V)-FLAG constructs. Cells were lysed and 60 μg of protein was used to measure triglyceride transfer activity in the presence or absence of the MTTP inhibitor lomitapide (1 μM)(% after 45 minutes); n = 9 (3 measurements from each of 3 independent experiments), mean +/-SD, One-way ANOVA with Bonferroni post-hoc tests, ** p<0.01, ***p<0.001). (G) Wild-type and mutant MTTP proteins were purified using anti-FLAG antibodies and used to measure phospholipid transfer in the presence or absence of lomitapide (1 μM) (180 minutes); n = 9 (3 measurements from each of 3 independent experiments), mean +/-SD, randomized block ANOVA with Bonferroni post-hoc tests, ** p<0.01, ***p<0.001.

The leucine residue mutated in the *stl* mutant fish (L475P) is conserved in human MTTP (L477) and lies within a highly conserved stretch of amino acids located in the middle alpha-helical domain of MTTP (Figure 6A). Analysis of mutations in patients with Abetalipoproteinemia has shown that missense mutations in the alpha-helical domain often prevent lipid transfer activity and result in loss of ApoB secretion (Reviewed in (Walsh and Hussain, 2017)). Therefore, it is likely that the *stl* L475P mutation alters the structure of the lipid-binding cavity in a similar manner to these patient mutations.

The glycine residue mutated in the C-terminus of *c655* mutants (G863V) is also conserved in the human sequence (G865) (Figure 6A). Unfortunately, it is unknown where the G865 residue is located in relation to the lipid-binding cavity because the C-terminal sequence of MTTP diverges from that of lipovitellin, so it is not modeled in the predicted MTTP structure. However, because the predicted tertiary structure of zebrafish Mttp is very similar to that of human MTTP and the sequence in the C-terminus is highly conserved (Figure 6A), we hypothesized that introduction of the *c655* mutation into the human MTTP protein would also cause loss of triglyceride transfer, but retention of phospholipid transfer activity. To test this hypothesis, we performed site-directed mutagenesis on the human *MTTP*-FLAG (hMTTP) plasmid and assessed the function of the MTTP (G865V) mutant protein.

COS-7 cells expressing the G865V mutant protein secreted slightly lower amounts (∼75%) of ApoB48 into the media compared to wild-type hMTTP, but this was significantly higher than the background levels seen in cells transfected with the empty vector (pcDNA3) (Figure 6B). ApoB48 concentrations in the cell lysate of cells expressing hMTTP and G865V mutant proteins were similar (Figure 6C). These data suggest that the G865V mutation supports ApoB48 secretion albeit at lower efficiency. Attempts were made to explain the mechanisms for lower efficiency in supporting ApoB secretion. Wild-type hMTTP-FLAG and G865V-FLAG were both localized to the ER (Fig. 6D), transfection efficiency was similar (36% hMTTP, 35% G865V), and the proteins were immunoprecipitated at similar levels from cell lysates (Figure 6E). Triglyceride transfer assays with cell lysates of COS-7 cells expressing hMTTP-FLAG and G865V-FLAG plasmids showed that the mutant G865V protein has significantly decreased (∼53%) triglyceride transfer activity compared to wild-type hMTTP, comparable to the activity level of the WT hMTTP in the presence the MTTP inhibitor lomitapide (1 μM) (Figure 6F). In contrast, the phospholipid transfer activity of hMTTP and G865V were not significantly different, and the activity of both alleles were inhibited to equal extent by lomitapide (Figure 6G). These data suggest that the *c655* mutation in both the zebrafish Mttp (G863V) and human MTTP protein (G865V) results in significant loss of triglyceride transfer activity, but has no effect on phospholipid transfer activity.

## DISCUSSION

The characterization of the zebrafish *mttp c655* mutation provides the first evidence that the triglyceride and phospholipid transfer functions of a vertebrate microsomal triglyceride transfer protein can be dissociated and identifies the putative region responsible for triglyceride transfer activity. Previous sequence comparisons of invertebrate and vertebrate orthologues of MTTP strongly suggested that acquisition of triglyceride transfer activity during evolution was the result of many changes in the lipid-binding cavity (Rava and Hussain, 2007), so it was unexpected that one missense mutation in the C-terminus selectively eliminated triglyceride transfer activity. Additionally, all of the characterized missense mutations from patients with Abetalipoproteinemia cause MTP to be either absent or deficient in both phospholipid and triglyceride transfer activities (Berthier et al., 2004; Di Filippo et al., 2012; Di Leo et al., 2005; Khatun et al., 2013; Miller et al., 2014; Narcisi et al., 1995; Rehberg et al., 1996; Ricci et al., 1995; Walsh et al., 2016; Walsh and Hussain, 2017; Walsh et al., 2015; Wang and Hegele, 2000).

While we do not know where the C-terminus is located in relation to the lipid-binding cavity (Figure 6A), one of the Abetalipoproteinemia mutations (G865X) results in a C-terminal truncation of 30 amino acids that prevents binding to PDI, thus resulting in loss of MTP protein (Ricci et al., 1995). Subsequent analysis of constructs *in vitro* indicated that deletion of the last 20 amino acids (Δ20, S875X), or mutating the cysteine at position 878 (C878S), reduced the expression of the protein to <15% of wild-type MTTP levels and abolished triglyceride transfer activity (Narcisi et al., 1995). This cysteine residue is conserved in MTTP orthologues from human to *C. elegans*, suggesting that it forms a disulfide bond essential for the tertiary structure of the protein. The authors argue that the residual protein produced must be binding to PDI, otherwise, no protein would be detected; however, it is unclear whether either of these mutated proteins are still capable of transferring phospholipid (Narcisi et al., 1995). In contrast to these mutations, the mutation of glycine to valine at position 865 in human MTTP (or 863 in zebrafish mttp) does not reduce protein expression or alter the localization of MTTP in the ER of COS-7 cells (Figure 5D, 6D), indicating that the mutation does not interfere with PDI binding. This suggests that the proposed disulfide bond is intact and that the G865V mutation may be more directly affecting lipid transfer. Our data show that this mutation lacks the ability to transfer triglycerides *in vitro*, but it is unclear whether the mutation also prevents binding of triglyceride in the lipid-binding cavity. Perhaps binding can still occur, which would be consistent with the evolutionary data, and maybe the C-terminus is necessary for transfer activity. Analyses of Drosophila, zebrafish, or human MTTP chimeric proteins may help test this hypothesis in the future. Additionally, in order to better understand how the *c655* mutation specifically inhibits triglyceride transfer, a crystal structure including the C-terminus will likely be necessary.

The production of B-lps in the ER of the intestine and liver is thought to occur in two steps. In the first step, MTP transfers lipids to ApoB as it is translated to form small primordial particles. In the second step, it has been suggested that fusion of ApoB-free lipid droplets in the lumen of the ER fuse to expand the lipoprotein core (“core expansion”) (Alexander et al., 1976; Boren et al., 1994; Hamilton et al., 1998; Wang et al., 1997). There is evidence to suggest that MTP is also responsible for producing these ER-lumenal lipid droplets (Kulinski et al., 2002). Using our LipoGlo assays, we have shown that the *c655* mutant fish produce small, homogenous, particles, whereas the wild-type embryos form VLDL-sized lipoproteins in the YSL at 2–3 dpf (Figure 4D). We have made similar observations in liver-specific *Mttp* KO mice expressing Drosophila *Mttp*, which has robust phospholipid transfer activity, but is deficient in triglyceride transfer (Khatun et al., 2012). Expression of fly Mttp resulted only in production of small B-lps, but human MTTP rescued the particle size (Khatun et al., 2012). Therefore, the phospholipid transfer activity of MTP may be crucial in the generation of the small homogenous particles representative of the first step of lipoprotein assembly, whereas triglyceride transfer might be primarily responsible for core expansion.

Although the effect of the *c655* mutation on the molecular function of the protein was unexpected, the lack of intestinal or hepatic steatosis is also consistent with our previous data with Drosophila Mttp expression in the livers of liver-specific Mttp-null mice. The phospholipid-rich high-density B-lps produced by the fly Mttp in hepatocytes partially restore plasma lipid levels and reduce liver steatosis (Khatun et al., 2012). Similarly, transfer of phospholipid and production of small B-lps in the *c655* mutant fish is not only sufficient for moving lipid from the liver, but is also capable of moving enough dietary lipid and fat-soluble vitamins from the intestine to prevent intestinal steatosis and support normal growth (Figure 3,4). Furthermore, retention of phospholipid transfer may also improve the health of the fish in ways that are independent of lipoprotein production. For example, MTP-dependent phospholipid transfer has been shown to be important for biogenesis and cell surface expression of CD1d and possibly other lipid-antigen-presenting molecules (Dougan et al., 2005).

In contrast to the health and viability of *c655* mutants, the *stl* mutant fish that survive have gross lipid accumulation in their intestine and severe growth defects (Figure 3,4), which is reminiscent of patients with Abetalipoproteinemia. The mutant protein localizes normally to the ER, but is deficient in triglyceride transfer, phospholipid transfer, and ApoB secretion (Figure 5). In the predicted model of the tertiary structure of Mttp, this mutation is located in the middle alpha-helical domain, not facing the lipid-binding cavity. The location and characterization of this mutation is very similar to that of the described Y528H and S590I Abetalipoproteinemia mutations, and it is likely that all three mutations prevent lipid transfer by altering the tertiary structure of the M subunit (Khatun et al., 2013; Miller et al., 2014).

Counter to the original characterization of *stl* mutants (Avraham-Davidi et al., 2012), we show that the *stl* mutants can survive to adulthood, especially when they are not competing with siblings for resources. We hypothesize that during the maintenance of this mutant line since its original characterization, a modifier has been eliminated that, when present in the *stl* background, was incompatible with life. In support of this hypothesis, the excessive sprouting angiogenesis defect for which the *stalactite* mutation was named (Avraham-Davidi et al., 2012), was also not as severe as originally described (Supplementary Figure S1). Whether the proposed modifier directly affects the secretion of B-lps, or some other aspect of development, is currently unclear. While the *stl* mutation severely decreases B-lp number and particle size (Figure 4D), some ApoB is still noted in the body of the *mttp^stl/stl^* embryos and larvae (Figure 4C, 4E). This suggests that there is enough transport of necessary lipids and fat-soluble vitamins by the small numbers of B-lps produced by the YSL and intestine to support development.

**Figure S1:**
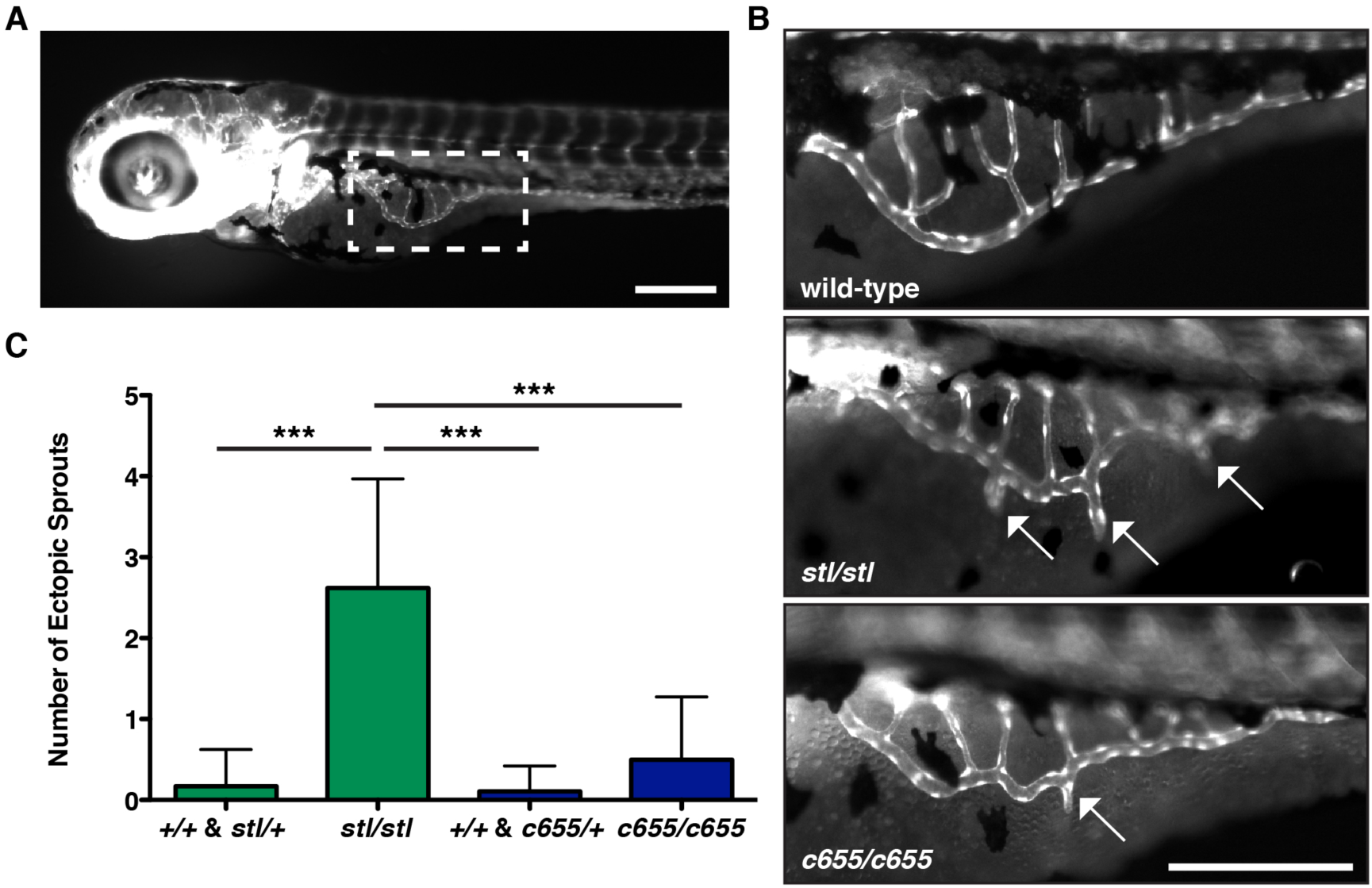
*c655*/*c655* embryos exhibit fewer ectopic angiogenic segments extending from the subintestinal vessels than *stl/stl* embryos. (A) The developing vasculature is visualized in the *Tg(fli:eGFP)^y1^* transgenic zebrafish line (Lawson and Weinstein, 2002). The subintestinal vessels (boxed region) grow bilaterally onto the dorsolateral surface of the yolk sac. Scale = 200 μM. (B) Representative wide-field images of *Tg(fli:eGFP)^y1^* in wild-type, *mttp^stl/stl^* or *mttp^c655/c655^* embryos at 3.5 dpf. Ectopic sprouts extending ventrally from the subintestinal vein are more common in *mttp^stl/stl^* mutant embryos than in *mttp^c655/c655^* mutant embryos. Scale = 200 μM. (C) Quantification of the average number of ectopic sprouts in *mttp* mutants and siblings on 3.5 dpf. Results represent pooled data from 3 independent experiments, n = 28-36 total embryos/genotype group; mean +/-SD, Kruskall-Wallis with Dunn’s Multiple Comparison test, *** p<0.001.

The *c655* mutants were initially identified due to the abnormal opaque appearance of their yolks, which is also a component of the *stl* mutant phenotype (Figure 1). We show that this opacity is due to the abnormal accumulation of lipid droplets in the yolk syncytial layer (Figure 2). As the lipids are liberated from the yolk, they are rapidly re-esterified to triglyceride and phospholipid and packaged into B-lps in the endoplasmic reticulum of the YSL. Because the *mttp* mutant fish are defective at producing (*stl*) and/or expanding (*c655*) B-lps, the re-esterified lipids are packaged instead into cytoplasmic lipid droplets. In contrast, cytoplasmic lipid droplets are only very rarely noted in the YSL of wild-type embryos, suggesting that the rate of yolk lipid break-down and the rate of B-lp production and secretion from the YSL are exquisitely coupled. We also noted that the size range of the cytoplasmic lipid droplets in the YSL of the *c655* mutants was much larger than in the *stl* mutants, with some LDs reaching 180 μm^2^. As the surface area-to-volume ratio decreases with increasing size spheres, the larger lipid droplets store more neutral lipid relative to the surface of the phospholipid coat. Because the *c655* mutants can transfer phospholipid to ApoB and secrete greater numbers of small dense B-lps than *stl* mutants, we hypothesize the retained neutral lipid is stored in larger lipid droplets because the phospholipid is being secreted and less available to coat lipid droplets. However, it is unclear whether the larger LDs form due to local production of triglycerides and cholesterol esters directly on their surface, or due to lipid droplet fusion, or both mechanisms (reviewed in (Olzmann and Carvalho, 2019; Walther et al., 2017)). The differences in the concentration and size of LDs between the mutants may result in differential effects on the degree of light scattering, which could explain the differences in opacity noted between mutants (Figure 1 – figure supplement 3).

None of the missense mutations identified in patients with Abetalipoproteinemia have been found to dissociate the lipid transfer activities of MTP (reviewed in (Walsh and Hussain, 2017)). However, given that the adult *c655* mutant zebrafish are indistinguishable from wild-type siblings, it is entirely possible that humans carrying a missense mutation that results in retention of phospholipid transfer exist in the population. While these people would likely have low plasma triglycerides, we would expect transport of fat-soluble vitamins to be normal. A thorough search of publicly available large human GWAS databases (Global BioBank Engine, T2D Knowledge Portal, GTEx Portal) did not reveal any coding variants near G865 other than the G865X mutation discussed above. However, since one copy of wild-type MTTP is sufficient to prevent fat malabsorption when faced with an oral fat load (Di Filippo et al., 2019), individuals heterozygous for a mutation similar to *c655* may not present with any changes in plasma lipid profiles.

Abnormally elevated levels of ApoB-containing lipoproteins and remnants promote atherosclerosis, the leading cause of death in the United States (CDC, 2018). Inhibition of MTP has long been considered a possible therapeutic target for lowering disease risk by inhibiting the production of VLDL and chylomicrons (Jamil et al., 1996; Wetterau et al., 1998)(for review see (Hussain and Bakillah, 2008; Walsh and Hussain, 2017)). Currently, the only MTP inhibitor approved for use in patients is lomitapide (Juxtapid®), which inhibits triglyceride and phospholipid transfer and reduces ApoB secretion (Robl et al., 2001)(Figure 6F,G). While this drug effectively reduces LDL cholesterol, total cholesterol, and plasma ApoB levels, it is only approved for patients with homozygous familial hypercholesterolemia, whose plasma cholesterol and triglyceride levels are up to four times the normal levels resulting in premature cardiovascular disease (Cuchel et al., 2014; Cuchel et al., 2013; FDA, 2012). While lomitapide effectively lowers circulating lipid levels and reduces cardiovascular disease risk in these patients, side effects include fat accumulation in the liver and adverse gastrointestinal events including reflux, indigestion, abdominal pain, constipation, and diarrhea (Blom et al., 2017; Cuchel et al., 2007; Cuchel et al., 2013).

The lack of intestinal and hepatic steatosis in the *c655* mutant fish suggests that an MTP inhibitor that selectively targets triglyceride transfer activity could potentially lower plasma lipids while preventing these gastrointestinal and liver side effects. This would not only improve the quality of life for patients currently taking lomitapide, but may also expand MTP inhibitor use to patients other than those with familial hypercholesterolemia. While one of the original MTP inhibitors discovered, BMS-200150, was very effective at inhibiting triglyceride transfer, but less effective (∼30%) at inhibiting phospholipid transfer in vitro (Jamil et al., 1996), later studies on purified MTP protein indicated the compound inhibits transfer of both lipid classes (Rava et al., 2006) and that it was not effective in animal models (Wetterau et al., 1998). Now that we appreciate that the triglyceride and phospholipid transfer functions of MTTP can be dissociated, we argue that it may be worth re-evaluating the phospholipid transfer activity of any previously identified compounds that inhibited triglyceride transfer activity of MTP, but failed to inhibit ApoB secretion *in vitro*. Perhaps new compounds could be specifically designed to target the C-terminal region of MTTP, although this will likely require a crystal structure of the MTP complex.

However, before a selective triglyceride transfer inhibitor could be considered as a therapeutic for a wide range of patients with hyperlipidemia, it will be important to determine whether the small lipoprotein particles produced by selective inhibition of triglyceride transfer are not atherogenic. Small dense LDL has been found to have greater potential for causing atherosclerosis than larger LDL sub-fractions and is a better predictor of cardiovascular disease than total LDL-cholesterol (Austin et al., 1988; Bjornheden et al., 1996; Ivanova et al., 2017). In future work, we want to evaluate whether the *mttp^c655/c655^* mutants are protected from atherosclerosis because they produce fewer B-lps or are at higher risk because the B-lp particles secreted are small and dense.

In conclusion, the unexpected discovery of the c655 missense mutation in *mttp* has provided novel insight into the structure-function relationship of MTP, underlining the importance of forward-genetic screening approaches to reveal aspects of biology that may otherwise be missed. Our work provides the first evidence that the triglyceride and phospholipid transfer functions of vertebrate MTP can be separated and that selective retention of phospholipid transfer is sufficient for dietary fat absorption and normal growth. These results argue that selective pharmacologic inhibition of triglyceride transfer may be a feasible therapeutic approach to treat disorders of lipid metabolism.

## Acknowledgments

We gratefully acknowledge Michael Sepanski for electron microscopy, Andrew Rock and Carmen Tull for fish husbandry, Matthew Bray for assistance using R, Amy Kowalski for synthesis of the pDESTTol2pA2-CMV: eGFP-CAAX plasmid, and Jennifer Anderson for help editing the manuscript. We would also like to especially thank Philip Ingham, who provided the *kif7* mutant strain in which we identified the *c655* mutation. This work was supported by NIH grants R01 DK093399 (Farber, PI; Busch-Nentwich, Co-PI), R01 GM63904 (The Zebrafish Functional Genomics Consortium; Ekker, PI, Farber, Co-PI), HL-137202-01A1 (Hussain, PI), R56 DK046900-17A1 (Hussain, PI) and F32DK109592 to M.H.W., as well as G. Harold & Leila Y. Mathers Foundation (Farber, PI), VA Merit Award BX004113-01A1 (Hussain, PI), AHA Postdoctoral Fellowship 19POST34410063 to S.R., and the Wellcome Trust [098051 and 206194] to E. Busch-Nentwich.

## Supplementary Files

Supplementary File 1: Single nucleotide variants present in *c655* mutant embryos.

## METHODS & MATERIALS

### Zebrafish husbandry and maintenance

Adult zebrafish were maintained at 27°C on a 14:10h light:dark cycle and fed once daily with ∼3.5% body weight Gemma Micro 500 (Skretting USA). Embryos were obtained by natural spawning and were raised in embryo medium at 28.5°C and kept on a 14:10 h light:dark cycle. All embryos used for experiments were obtained from pair-wise crosses and were staged according to (Kimmel et al., 1995). Exogenous food was provided starting at 5.5 days post fertilization (dpf) unless otherwise noted. Larvae were fed with GEMMA Micro 75 (Skretting) 3x a day until 14 dpf, GEMMA Micro 150 3x a day + Artemia 1x daily from 15 dpf–42 dpf and then GEMMA Micro 500 daily supplemented once a week with Artemia. Zebrafish sex is not determined until the juvenile stage, so gender is not a variable in experiments with embryos and larvae. Sex of adult fish included in analyses is noted in Figure legends. All zebrafish protocols were approved by the Carnegie Institution Department of Embryology Animal Care and Use Committee (Protocol #139).

*Stalactite* (*stl*) *mttp* mutant zebrafish in the *Tg(fli1:eGFP)^y1^* background (Avraham-Davidi et al., 2012; Lawson and Weinstein, 2002; Yaniv et al., 2006) were provided by Karina Yaniv (Weizmann Institute of Science, Israel) and out-crossed to the AB wild-type strain. The *stl* mutation was maintained in both the presence and absence of the *fli1:eGFP* transgene. The *c655* phenotype was identified in the Farber laboratory in the background of a *kif7* mutant strain that was obtained from Philip Ingham (Lee Kong Chian School of Medicine, Singapore). The *c655 mttp* mutation was isolated from the *kif7* mutation by out-crossing to the AB wild-type strain. The *c655* mutation was crossed into the *Tg(fli1:eGFP)^Y1^* reporter line. Both *stl* and *c655 mttp* mutations were crossed into the *ApoBb.1-NanoLuc* LipoGlo reporter line (Thierer et al., In Press).

### Positional Cloning

To map the location of the mutation responsible for the c655 phenotype, 23 embryos with normal yolks and 23 embryos with opaque yolks (3 dpf) were processed for RNA-seq (White et al., 2017). RNA was extracted from embryos by mechanical lysis in RLT buffer (Qiagen, 79216) containing 1 μL of 14.3M beta-mercaptoethanol (Sigma, M6250). The lysate was combined with 1.8 volumes of Agencourt RNAClean XP (Beckman Coulter, A63987) beads and allowed to bind for 10 minutes. The plate was applied to a plate magnet (Invitrogen) until the solution cleared and the supernatant was removed without disturbing the beads. This was followed by washing the beads three times with 70% ethanol. After the last wash, the pellet was allowed to air dry for 10 mins and then resuspended in 50 μl of RNAse-free water. RNA was eluted from the beads by applying the plate to the magnetic rack. RNA was quantified using the Quant-iT 610 RNA assay (Invitrogen, Q33140). Total RNA from individual embryos was DNase treated for 20 mins at 37°C followed by addition of 1 μL 0.5M EDTA and inactivation at 75°C for 10 mins to remove residual DNA. RNA was then cleaned using 2 volumes of Agencourt RNAClean XP (Beckman Coulter, A63987) beads under the standard protocol. Strand-specific RNA-seq libraries containing unique index sequences in the adapter were generated simultaneously following the dUTP method using 700 ng total RNA and ERCC spike mix 2 (Ambion, 4456740). Libraries were pooled and sequenced on Illumina HiSeq 2500 in 75bp paired-end mode. Sequence data were deposited in European Nucleotide Archive under accession ERP023267. FASTQ files were aligned to the GRCz10 reference genome using TopHat2 (Kim et al., 2013) (v2.0.13, options: --library-type fr-firststrand). Ensembl 88 gene models were supplied to TopHat2 to aid transcriptome mapping. MMAPPR (Hill et al., 2013) was used to determine the location of the causal mutation. Variants were called from the pooled data using the GATK HaplotypeCaller (Van der Auwera et al., 2013). Variants inside the regions output by MMAPPR were selected and filtered for ones where the mutant sample was called as being homozygous alternate and the siblings were heterozygous. The consequences of these variants on annotated genes was calculated using the Ensembl Variant Effect Predictor (McLaren et al., 2016) and SIFT (Sim et al., 2012). Variants with the following consequences were selected as candidates for the causal mutation: stop_gained, splice_donor_variant, splice_acceptor_variant, transcript_ablation, frameshift_variant, stop_lost, initiator_codon_variant, missense_variant, inframe_insertion, inframe_deletion, transcript_amplification, splice_region_variant, incomplete_terminal_codon_variant.

### DNA Extraction and Genotyping

Genomic DNA was extracted from embryos or adult fin clips using a modified version of the HotSHOT DNA extraction protocol (Meeker et al., 2007). Embryos/tissues were heated to 95°C for 18 minutes in 100 μL of 50 mM NaOH. The solution was cooled to 25°C and neutralized with 10 μL of 1M Tris-HCl pH 8.0. Genotyping primers for the stalactite allele were designed using the dCAPS Finder 2.0 program (Neff et al., 2002) and synthesized by Eurofins Genomics. The *stalactite* locus was amplified using the forward primer 5’-GTC TGA GGT TCA GAT GTA CCT GTT AGG AC-3’ and reverse primer 5’-CTC TGC TGT GAT GAG CGC AGG-3’ (0.5 μM primer, T_a_ = 60°C, extension time 30”). The forward primer introduces an AvaII restriction site into the mutant amplicon, such that following digestion (5 units of AvaII (New England BioLabs, R0153) at 37°C, 4 h) the WT band is 157bp, homozygous mutants have bands at 129 bp and 28 bp, and heterozygotes have all three bands. The *c655* locus was amplified using the forward primer 5’-AGAGACGGTGTCCAAGCAGG-3’ and reverse primer 5’-GCTCAAAGACTTTCTTGC-3’ (0.25 μM primer, Ta = 50°C, extension time 30”). The *c655* mutation introduces a BsrI restriction site into the amplicon, such that following digestion (3 units of BsrI (New England BioLabs, R0527) in NEB Buffer 3.1 (B7203), 65°C, 3.5 h) the WT band is 137 bp, homozygous mutants have bands at 76 bp and 61 bp, and heterozygotes have all three bands. For the *ApoBb.1-NanoLuc* genotyping protocol, see (Thierer et al., In Press).

### Generation of mttp-FLAG and ApoB48 plasmids

The wild-type zebrafish *mttp* coding sequence with a FLAG-tag prior to the termination codon at the C-terminus was generated by custom gene synthesis and cloned into the pcDNA3.1+ vector (*mttp*-FLAG)(Gene Universal Inc., Newark, DE). The *stl* and *c655* mutations were subsequently introduced to this plasmid by site-directed mutagenesis (Gene Universal Inc.) to generate *mttp^stl^*-FLAG and *mttp^c655^*-FLAG plasmids. The human pcDNA3.1-*MTTP*-FLAG plasmid was synthesized as described previously (Rava et al., 2006; Sellers et al., 2003). The human equivalent of the *c655* mutation (G865V) was introduced into this plasmid using the Q5 Site-directed mutagenesis kit (New England Biolabs), with the following primer pair: Forward 5’-CGTATTAGCAgtaTGTGAATTCC-3’, Reverse 5- CTTTCTTTTCTTTTCTGAGAG-3’. The human ApoB48 sequence (Hussain et al., 1995) was cloned into the pcDNA3 under control of the CMV promoter.

### Rescue of c655 opaque yolk phenotype

*mttp^c655/c655^* embryos were injected at the 1-cell stage with 20pg of zebrafish *mttp*-FLAG plasmid and 20 pg of CMV:*eGFP-CAAX* (synthesized using the Tol2kit Gateway cloning system using the p5E-CMV/SP6, pME-*eGFP-CAAX*, and p3E-polyA entry clones (Kwan et al., 2007)) as a marker of successful injections. Embryos were raised to 3 dpf and screened for eGFP expression in the yolk sac. Images of eGFP+ control and experimental embryos were blinded and scored for yolk opacity by another member of the lab.

### Ectopic sprout analysis

*mttp^stl/stl^*, *mttp^c655/c655^* and WT zebrafish in the *Tg(fli1:eGFP)^y1^* background were imaged at 3 dpf with a Zeiss Axiozoom V16 microscope equipped with a Zeiss PlanNeoFluar Z 1x/0.25 FWD 56mm objective, AxioCam MRm camera, and Zen 2.5 software. The length of ectopic angiogenic segments that extend from the subintestinal vessels were analyzed in Fiji (Schindelin et al., 2012)(ImageJ V2.0.0, National Institutes of Health, USA) as described by (Avraham-Davidi et al., 2012).

### Transmission Electron Microscopy

Wild-type, *mttp^stl/stl^*, *mttp^c655/c655^,* and *mttp^stl/c655^* mutant zebrafish embryos were fixed at 4 dpf in a 3% glutaraldehyde, 1% formaldehyde, 0.1 M cacodylate solution for 1-3 h. Embryos were trimmed and swim bladders were deflated before embedding in 2% low melt agarose and processed as described in (Zeituni et al., 2016). Post-fixation was performed for 1 h with 1% osmium tetroxide + 1.25% potassium ferricyanide in cacodylate solution. Following 2 x 10 min washes with water, samples were incubated with 0.05M maleate pH 6.5 for 10 min. Samples were stained en bloc with 0.5% uranyl acetate in maleate for 4°C overnight. Following 2 x 15 min washes with water, samples were dehydrated through graded EtOH dilution (35%, 2 x 15 min; 50%, 15 min; 75%, 15 min; 95%, 15 min; 100% 4 x 15 min). Samples were washed with propylene oxide 4 x 15 min before incubation with 1:1 propylene oxide/resin (Epon 812 epoxy, Ladd Research Industries, Williston, VT) for 1 h and evaporated overnight. This was followed by 2 x 1 h washes in 100% resin and a final embedding in 100% resin at 55°C overnight followed by 70°C for three days. Sections were made on a Reichert Ultracut-S (Leica Microsystems), mounted on naked 200 thin mesh grids, and stained with lead citrate. Images were obtained with a Phillips Technai-12 electron microscope (FEI, Hillsboro, OR) and 794 Gatan multiscan CCD camera (Gatan, Pleasanton, CA) using Digital Micrograph software. Lipid droplet number and area was quantified with Fiji.

### Growth Time-Course

Unsorted embryos from pair-wise in-crosses of *stalactite* or *c655* heterozygous fish and pair-wise crosses of *mttp^stl/+^* x *mttp^c655/+^* were raised and were analyzed for standard length at 1, 3, 6, 9, 12, and 24 weeks post fertilization. At 1 week, fish were imaged using a Nikon SMZ1500 microscope with HR Plan Apo 1x WD 54 objective, Infinity 3 Lumenera camera and Infinity Analyze 6.5 software. Standard length (Parichy et al., 2009) was measured using Fiji (NIH). Starting at three weeks, standard length was measured with a ruler. Mass of the fish was also measured starting at 6 weeks. At 1 and 3 weeks, gDNA was obtained from whole fish for genotyping. At later time-points, genotyping was performed on fin clips. Images of fish at 12 weeks post fertilization were taken with a Canon T6 camera with a Canon EF 100mm Macro Lens.

### Tissue Histology

Adult zebrafish (7.5 mo; 2 males, 1 female per genotype) were placed individually into mating tanks and fasted overnight (∼24 h). Fish were euthanized by submersion in ice-water. A piece of the anterior intestine and the liver were dissected from each animal and fixed in neutral-buffered formalin (Sigma, F8775) at 4°C for 48 h. Sectioning and hematoxylin & eosin staining was performed by the Johns Hopkins University Oncology Tissue Services. Slides were imaged with a Nikon E800 microscope with 60×/1.4 oil Plan Apo Nikon objective and Canon EOS T3 camera using EOS Utility image acquisition software.

### Tissue Lipid Extractions, HPLC & Analysis

Adult zebrafish (1 yr; 2–3 males, 2–3 females per genotype) were fasted overnight (∼24 h) and euthanized by submersion in ice-water. Similar size pieces of the anterior intestine and the liver were dissected from each animal and frozen on dry ice. Tissues were sonicated in 550 μL of homogenization buffer (20 mM Tris-HCl, 1 mM EDTA), and the protein concentration of each sample was measured using the BCA protein assay kit (Pierce, 23225). Lipids were extracted from the remaining sample volume by a modified Bligh-Dyer procedure (Carten et al., 2011), dried under vacuum, and re-suspended in 50 μL of HPLC-grade isopropanol as the HPLC injection solvent. Injection volumes were optimized for each sample to produce peak shapes appropriate for quantitation (1–25 μL). The lipid components of each sample were separated and detected by an HPLC-CAD system using a LPG-3400RS quaternary pump, WPS-3000TRS autosampler (maintained at 20°C), TCC-3000RS column oven (maintained at 40°C), Accucore C18 column (150 x 3.0 mm, 2.6 μm particle size), FLD-3100 fluorescence detector (8 mL flow cell maintained at 45°C), and a Dionex Corona Veo charged aerosol detector (Thermo Fisher Scientific). Lipids were separated over an 80 min time range in a multi-step mobile phase gradient as described in (Quinlivan et al., 2017). The lipid class and area of each analyte peak were determined using Chromeleon 7.2 (Thermo-Fisher Scientific) for chromatogram visualization and manual integration as described in (Otis et al., 2017). For quantitative comparisons between samples, each lipid peak area was normalized to the protein concentration of the homogenized tissue.

### LipoGlo Assays

All LipoGlo assays were performed with fish carrying a single copy of the LipoGlo *(apoBb.1^NLuc/+^)* reporter. For detailed LipoGlo methods see (Thierer et al., In Press); Nano-Glo reporter system reagents are all from Promega Corp., (N1110; (Hall et al., 2012)). For quantitative assays and B-lp size analysis, individual embryos were dispensed into 96-well plates (USAScientific, #1402-9589) and homogenized in 100 μL of B-lp stabilization buffer (40 mM EGTA, pH 8.0, 20% sucrose + cOmplete mini, EDTA-free protease inhibitor (Sigma, 11836170001)) by sonication in a microplate-horn sonicator (Qsonica Q700 sonicator with a Misonix CL-334 microplate horn assembly). Homogenate was stored on ice for immediate use or frozen at −20°C for later use. ApoB-Nanoluc levels were quantified by mixing 40 μL of embryo homogenate with an equal volume of diluted Nanoluc buffer (1:3 NanoGlo buffer:PBS + 0.5% NanoLuc substrate (furimazine)) in a 96-well opaque white OptiPlate (Perkin-Elmer, 6005290), and the plate was read within 2 minutes of buffer addition using a SpectraMax M5 plate reader (Molecular Devices) set to top-read chemiluminescent detection with a 500 ms integration time. To quantify the size distribution of B-lps, 12 μL of homogenate was combined with 3 mL of 5x loading dye (40% sucrose, 0.25% bromophenol blue, in Tris/Borate/EDTA (TBE) buffer), and 12.5 mL of the resulting solution (10% larval homogenate) was loaded per well on a 3% native polyacrylamide gel. Each gel included a migration standard of Di-I-labeled human LDL (L3482, ThermoFisher Scientific). Gels were run at 50V for 30 min, followed by 125V for 2 h. Following application of 1 mL of TBE supplemented with 2 μL of Nano-Glo substrate to the surface of the gel and incubating for 5 min, gels were imaged with an Odyssey Fc (LI-COR Biosciences) gel imaging system. Images were obtained in the chemiluminescent channel (2 min exposure) and then the 600 nm channel (30 sec) for Nanoluc detection and Di-I LDL standard detection, respectively. Each lane on the gel was converted to a plot profile in Fiji and divided into LDL, IDL, VLDL and Zero Mobility bins based on migration relative to the Di-I LDL standard. Pixel intensity from the plot profile was summed within each bin for comparison between genotypes. To determine the localization of B-lps in the whole fish, intact embryos or larvae were anesthetized and fixed in 4% paraformaldehyde for 3 h at room temperature. Following rinses in PBS + 0.1% tween-20 (3 x 15 min), embryos were mounted in 1% low-melt agarose (BP160-100, Fisher Scientific) in TBE supplemented with 1% Nano-Glo substrate. Chemiluminescent images (10 and 30 sec exposures with no illumination) and a brightfield image were taken with a Zeiss Axiozoom V16 microscope equipped with a Zeiss Plan NeoFluar Z 1x/0.25 FWD 56mm objective, AxioCam MRm camera, and Zen 2.5 software, using 2×2 binning and 2x gain. Images were quantified using Fiji; regions of interest (ROI) were drawn on the brightfield image (viscera, trunk, and head), and these ROIs were used to quantify the NanoLuc intensity on the 30sec exposure chemiluminescent images. ROIs of the same shape were used to calculate the background signal, which was subtracted from the intensity value for each ROI.

### Oil Red O staining

15-dpf larval zebrafish were fixed with 4% paraformaldehyde in PBS for 3 h at room temperature and then overnight at 4°C. Fish were rinsed in 60% 2-propanol for 10 minutes, rocking and then put into 0.3% Oil Red O (Sigma-Aldrich, #O0625) to rock overnight at room temperature. Fish were rinsed 3 times with 60% 2-propanol for 15 minutes. Washed fish were equilibrated step-wise into glycerol and imaged with incident light using a Nikon SMZ1500 microscope with HR Plan Apo 1x WD 54 objective, Infinity 3 Lumenera camera, and Infinity Analyze 6.5 software.

### ApoB secretion assays

Monkey kidney COS-7 cells which do not express MTTP or ApoB were plated in 10 cm^2^ cell culture dishes at a density of 9 x 10^5^ cells per plate and grown in Dulbecco’s modified Eagle’s medium (DMEM) containing 10% fetal bovine serum, L-glutamine, and antibiotics at 37°C. COS-7 cells were transfected with 5 μg of plasmid expressing human ApoB48 cDNA under the control of CMV promoter using endofectin (Genecopoeia, EF014) according to the manufacturer’s protocol. After 24 hours, cells from each dish were harvested, equally distributed in 6-well plates, and reverse transfected with 3μg of either pcDNA3, pcDNA3-*mttp*-FLAG, pcDNA3-*mttp^stl^*-FLAG, pcDNA3-*mttp^c655^*-FLAG, pcDNA3-*MTTP*-FLAG, or pcDNA3-*MTTP*(G865V)-FLAG plasmids. After 32 h cells were incubated overnight with 1 mL of DMEM containing 10%FBS. The overnight conditioned media were collected to measure ApoB by ELISA (Bakillah et al., 1997; Hussain et al., 1995). Cells were scraped in PBS and a small aliquot was used to measure total protein using a Coomassie protein assay (Thermo Scientific, #1856209). Cells were lysed in cell extract buffer (100 mM Tris, pH 7.4, 150 mM NaCl, 1 mM EGTA, 1mM EDTA, 1% Triton X-100, 0.5% sodium deoxycholate). Lysates were rotated for 1 h at 4 °C to solubilize the membranes and centrifuged at 16,000g for 30 mins. ApoB was measured in the supernatant via ELISA. Briefly, high binding 96 well plates (Corning, #3366) were incubated with capture antibody anti-LDL (apoB), clone 1D1(MyBiosource, #MBS465020, 1:1000 dilution) overnight at room temperature. The plate was washed 3x with PBS-T (PBS + 0.05% Tween-20) and blocked with 3% BSA (Boston Bio Products, #P753) for 1 h and washed 3x with PBS-T, before incubating with 100 μL of standards and experimental samples for 3 h. The plate was washed 3x with PBST and incubated with 100 μL of human ApoB antibody (Academy Bio-Medical Company, Inc., #20S-G2, 1:1000 dilution) for 1 h. After washing the plate 3x with PBS-T, 100 μL of alkaline phosphatase labeled anti-goat IgG (Southern Biotech, #6300-04, 1:3000 dilution) was added to each well and incubated for 1 h. The plate was washed 3x with Diethanolamine buffer, pH 9.5 and 100 μl of PNPP (Thermo Scientific, #34045, 1 mg/mL) was added to each well before reading the plate at 405nm in a PerkinElmer Victor^3^ 1420 multilabel counter. Data for zebrafish and human plasmids were obtained in the same experiments, but are graphed separately in figures 5A,B & 6B,C; the pcDNA control data is displayed in both sets of graphs.

### Immunofluorescence

COS-7 cells were plated at a density of 50,000 cells on coverslips in 12-well dishes and transfected with 2 μg of plasmids expressing either zebrafish or human MTTP-FLAG plasmids. After 48 h, cells were fixed in paraformaldehyde and blocked with PBS supplemented with 1 mM MgCl_2_, 0.5 mM CaCl_2_, 3% BSA, 0.1% Triton X-100 and 1% horse serum. Cells were incubated with anti-FLAG M2 monoclonal antibody (Sigma # F3165, 1:250 dilution) and anti-calnexin antibody (Santa Cruz Biotechnology, # sc-11397, 1:250 dilution) overnight. Cells were washed three times with PBS and incubated with goat anti-mouse Alexa Fluor-594 (Invitrogen, #A11005, 1:500 dilution) and donkey anti-rabbit Alexa-Fluor-488 (Invitrogen, # A21206, 1:500 dilution) for 1 h. The cells were washed and mounted with Vectashield mounting medium (Vector Laboratories, #H-1000). Images were taken on a Leica SP5II confocal microscope with a 63×1.4 HCX PL Apo oil immersion lens.

### Immunoprecipitation and Western blotting

Transfected COS-7 cells were washed three times with ice cold PBS and scraped in buffer K (1 mM Tris-HCl, 1 mM EGTA and 1 mM MgCl_2_, pH 7.6) containing protease inhibitor cocktail (Sigma, # P2714). Cells were mechanically lysed by passing them 10 times with 30_1/2_-gauge needle and small fractions were used to measure the total protein using a Coomassie protein assay (Thermo Fisher Scientific, #1856209). Cell lysate was incubated with Anti-FLAG M2 antibody for 1 h and immunoprecipitated (IP) using (protein A/G) agarose beads (Santa-Cruz Biotechnology, # SC2003). The supernatants were used to detect actin via Western blotting and served as loading controls. Both the supernatant and immunoprecipitated fractions were subjected to electrophoresis on an 8% SDS-PAGE gel. The weight separated proteins were transferred to nitrocellulose membranes and probed with either anti-FLAG M2 (1:1000) or anti-actin (Thermo Fisher Scientific, #PA1-183, (1:3000)) prepared in 2% BSA in TBS. The blots were washed and probed with HRP-conjugated corresponding secondary antibodies (goat anti-rabbit, Cell Signaling Technology, #7074, 1:5000 or goat anti-mouse, Thermo Fisher Scientific, #62-6520, 1:5000). The blots were developed in ChemiDoc^TM^-Touch Imaging system from Bio Rad.

### Triglyceride transfer assay

Following transfection with plasmids as described above, cell lysate (35 μg) prepared in buffer K containing protease inhibitor cocktail was incubated with donor vesicles containing NBD-labeled triolein (Setareh Biotech, LLC, #6285) and acceptor vesicles. Fluorescence was measured at different time intervals (5, 10, 15, 30, 45 and 60 mins). Percent triglyceride transfer was calculated after subtracting the blank and dividing it by the total fluorescence reading obtained by disrupting vesicles with isopropanol, as described previously (Athar et al., 2004; Rava et al., 2005). Where noted, assays also included the MTTP inhibitor Lomitapide (Aegerion Pharmaceuticals, #AEGR-733) at a concentration of 1 μM.

### Phospholipid transfer assay

COS-7 cells were transfected with 9 μg of either zebrafish *mttp*-FLAG or human MTTP-FLAG plasmids in 10 cm^2^ cell culture dishes. After 48 h, cell lysates were prepared in buffer K containing protease inhibitor cocktail (Sigma, #P2714). The cell lysates were centrifuged at 12,000g for 10 mins at 4°C. A small aliquot of cell lysate was used for measuring protein and kept for Western blotting to measure expression level. Equal concentrations of protein from each sample were incubated with 40 μL of M2 agarose beads (Sigma, #A2220) for 3 h at 4°C. FLAG-tagged protein were eluted twice with 200 ng/μL FLAG peptide (Sigma, #F3290; 1 h at 4°C). PL transfer activity was assayed using NBD labeled Phosphoethanolamine, triethylammonium (Thermo Fisher Scientific, #N360). The purified FLAG-tagged proteins were incubated with donor vesicles containing NBD-Phosphoethanolamine and acceptor vesicles. The fluorescence was measured at different time intervals (1, 2, 3 and 4 h). The percentage transfer of phospholipid was calculated as the difference between the fluorescence reading at the 0 h time point and 3 h time point divided by the total fluorescence reading obtained by disrupting vesicles with isopropanol as described previously (Athar et al., 2004; Rava et al., 2005). Where noted, assays also included the MTTP inhibitor Lomitapide at a concentration of 1 μM.

### Modeling

Predicted models of human MTTP and zebrafish Mttp proteins were generated using Phyre2 (Kelley et al., 2015) and the graphics were enhanced for clarity using Chimera (UCSF Chimera, developed by the Resource for Biocomputing, Visualization, and Informatics at the University of California, San Francisco, with support from NIH P41-GM103311 (Pettersen et al., 2004).

### Statistical Analyses

Graphing and some statistics, including One-way, randomized block and Repeated Measures ANOVA with Bonferroni post-hoc tests, Kruskall-Wallis with Dunn’s Multiple Comparison test and Chi-square tests were performed with GraphPad Prism (GraphPad Software). When sample sizes and variance between groups were significantly different, Robust ANOVA was performed using R to determine overall significance of noted datasets (Mair and Wilcox, 2018)(https://cran.r-project.org/web/packages/WRS2/vignettes/WRS2.pdf),((Mangiafico, 2015), https://rcompanion.org/rcompanion/d_08a.html). When significant differences were present between genotypes, Games-Howell post-hoc tests were used to make pair-wise comparisons at each time point using SPSS Statistics (IBM), adjusting the significance level for multiple comparisons. Details of the statistical analyses can be found either in the figure legend or results sections. Sample sizes for each experiment are indicated in the figure legends for each experiment.

### Additional Software

DNA, mRNA, and protein sequence alignments were performed with MacVector V15.5 (MacVector, Inc.). Microsoft Word and Excel were used for manuscript preparation and data analysis, respectively, figures were assembled in Adobe Illustrator CS5 (Adobe Systems) and references were assembled with EndNote 8X.

